# Optimizing 5’UTRs for mRNA-delivered gene editing using deep learning

**DOI:** 10.1101/2023.06.15.545194

**Authors:** Sebastian Castillo Hair, Stephen Fedak, Ban Wang, Johannes Linder, Kyle Havens, Michael Certo, Georg Seelig

## Abstract

mRNA therapeutics are revolutionizing the pharmaceutical industry, but methods to optimize the primary sequence for increased expression are still lacking. Here, we design 5’UTRs for efficient mRNA translation using deep learning. We perform polysome profiling of fully or partially randomized 5’UTR libraries in three cell types and find that UTR performance is highly correlated across cell types. We train models on all our datasets and use them to guide the design of high-performing 5’UTRs using gradient descent and generative neural networks. We experimentally test designed 5’UTRs with mRNA encoding megaTALTM gene editing enzymes for two different gene targets and in two different cell lines. We find that the designed 5’UTRs support strong gene editing activity. Editing efficiency is correlated between cell types and gene targets, although the best performing UTR was specific to one cargo and cell type. Our results highlight the potential of model-based sequence design for mRNA therapeutics.

## Introduction

mRNA therapeutics and vaccines provide a safe, effective, and flexible method of delivering transient genetic instructions to living cells and tissues^1^. Compared to plasmid or AAV-based delivery, mRNA offers several advantages, including simple manufacturing that is independent of the encoded therapeutic protein^2^, lower immunogenicity, and transient gene expression^3,4^. As a result, mRNA technology has been crucial for the rapid development of vaccines against the COVID-19 pandemic^5,6^, and is currently being developed for applications such as protein replacement therapy^7,8^, regenerative medicine^9,10^, and cancer immunotherapy^11,12^, among others^13^. An intriguing use of the mRNA platform is delivery of gene editing reagents^3,14^, because transient expression of gene editors avoids deleterious effects from prolonged exposure such as off-target editing^4^ and reduces the likelihood of forming anti-drug antibodies, thereby allowing for repeated dosing^15^. Though there are multiple gene editing platforms, single-chain compact enzymes such as megaTALs^16^ are particularly well-suited to mRNA delivery. megaTALs are fusions of a minimal transcription activator-like (TAL) effector domain with an engineered meganuclease. The TAL effector addresses the meganuclease, which has intrinsic specificity for a few genomic target sites, to a single site where it catalyzes the formation of a DNA double stranded break, thereby achieving high activity and specificity^16^. Because of these features, megaTALs have been developed for a number of therapeutically-relevant targets^17–19^.

The recent success of mRNA vaccines and therapeutics is the result of decades of research in areas such as lipid nanoparticles for delivery^20^, modified nucleosides for decreased immunogenicity^21,22^, 5’-cap analogs for improved translation and stability^23^, and codon optimization^24^. Comparably little attention has been directed to optimizing untranslated regions (UTRs) despite their roles in controlling mRNA translation and stability. Many mRNA therapies currently utilize UTRs from the alpha- and beta-globin genes or slight modifications thereof, owing to being well described and associated with highly expressed proteins^1,13^. However, increased protein expression has been observed in a few studies when using alternative UTRs^25–28^, demonstrating the remaining untapped potential to optimize expression. A major obstacle is the difficulty in predicting the effects of arbitrary UTR sequences, as some cis-regulatory elements can affect multiple molecular processes^29^ and interact with RNA-binding proteins^30^ and microRNAs^31,32^ that may even be differentially expressed across tissues. Recently, quantitative models based on deep learning that predict translation efficiency^33,34^ and mRNA stability^35^ from sequence have started to emerge. Using these to guide UTR sequence design for mRNA therapeutics remains an intriguing yet relatively unexplored alternative^36^ (**Figure 1A**).

**Figure 1.**
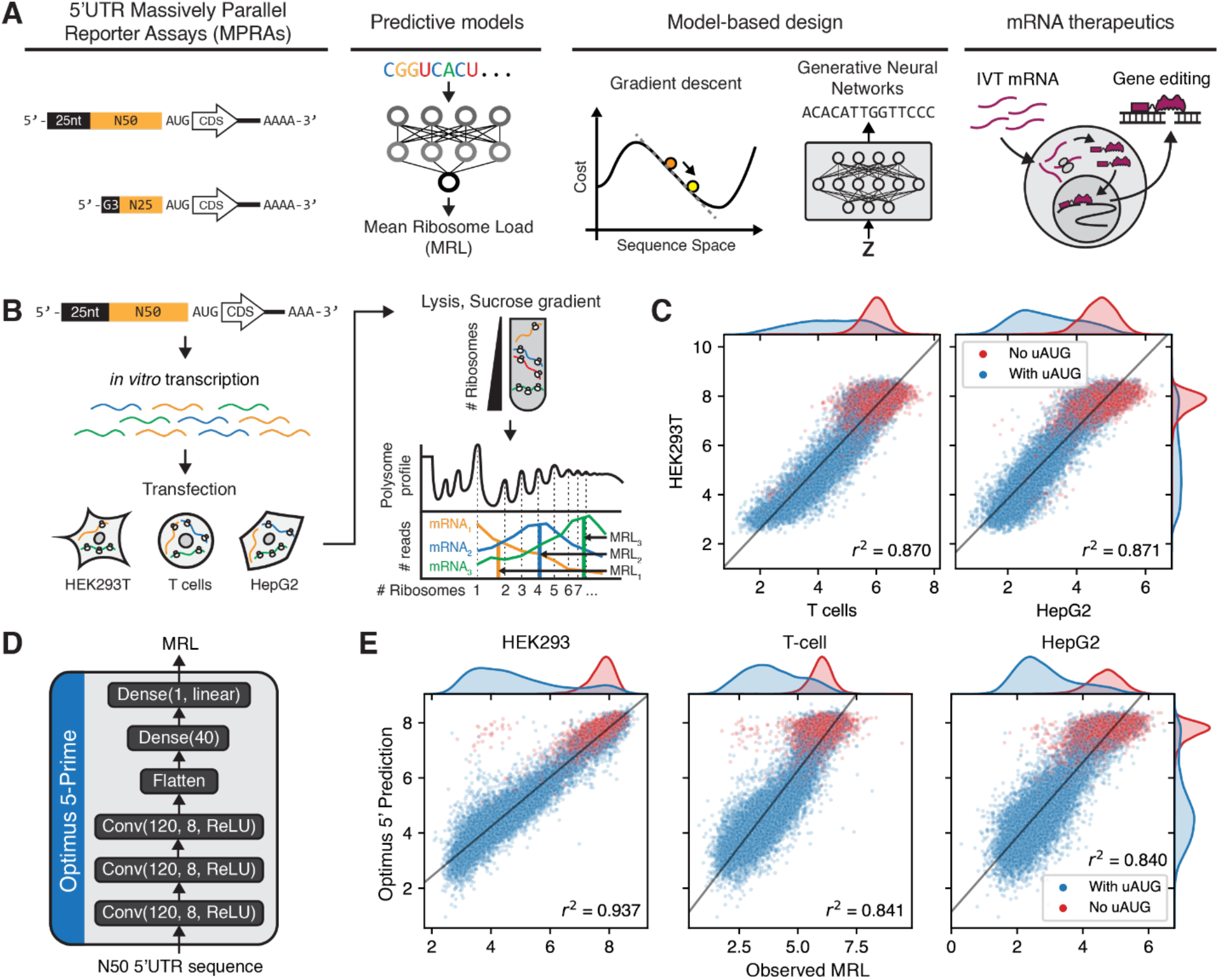
Massively Parallel Reporter Assays (MPRAs) to measure cell type-specific 5’UTR regulation of translation. **(A)** A model-based design strategy for 5’UTRs in mRNA therapeutics applications, using neural network-based predictive models trained on MRPA data. **(B)** Summary of polysome profiling MPRA. A library with a randomized 50nt 5’UTR region was synthesized as *in vitro* transcribed (IVT) mRNA, transfected into HEK293T, T cells, and HepG2 cells, and fractionated using a sucrose gradient to separate mRNAs with different numbers of ribosomes. Fractions were then barcoded and sequenced, and the Mean Ribosome Load (MRL) was calculated for each 5’UTR variant as a proxy of translation efficiency. The resulting data contained 204,803 5’UTR variants with 100 or more reads in all replicates, in two replicates in HEK293T, two in T cells, and one in HepG2. **(C)** Comparison of MRL measurements across cell lines. 5’UTR variants were sorted by the minimum number of reads across all replicates, and the top 20,000 were used for this analysis. **(D)** Architecture of Optimus 5-Prime, a convolutional neural network model for predicting MRL from 5’UTR sequence^33^. **(E)** Optimus 5-Prime predictions compared to MRL measurements in all three cell lines. The top 20,000 5’UTRs by read count in HEK293, which were not used for model training, were used for this analysis.

The 5’UTR sequence in particular is a major determinant of translation efficiency and thus an intriguing target for engineering^37–39^. To initiate translation, the ribosomal 43S pre-initiation complex (PIC) scans the 5’UTR in the 5’-to-3’ direction until a start codon is found. Therefore, 5’UTRs can affect translation by capturing PICs prematurely via upstream start codons (uAUGs) and ORFs (uORFs)^38,40^, interfering with PIC scanning via stable secondary structure^39^, or even directly recruiting ribosomes via Internal Ribosome Entry Sites (IRESs)^41^. Some 5’UTR cis-regulatory elements are exclusively located within a few bases from the 5’ end. For example, 5’-Terminal Oligo Pyrimidine (5’TOP) motifs consisting of a cytosine at position +1 followed by 4 to 15 pyrimidines^42^, 5’TOP-like motifs located within four nucleotides of the transcription start site^43^, and pyrimidine-rich translational elements (PRTEs) consisting of a uridine flanked by pyrimidines^44,45^ upregulate translation in response to mTOR activation during stress and have been linked to cancer initiation and progression. Transcriptome-wide translation measurements in a panel of cell lines^46^ and during neuronal differentiation^47^ have suggested that 5’UTRs regulate translation in a mostly cell type-independent manner whereas 3’UTRs have a greater cell type-specific effect. However, some 5’UTRs have been observed to act in a cell type-specific manner, for example during embryo development^48,49^.To predict the influence of 5’UTR sequence on translation, we previously developed Optimus 5-Prime, a convolutional neural network trained on translation efficiency measurements from a synthetic reporter library of 280,000 random 5’UTRs^33^. However, there are limitations to Optimus 5 Prime. First, the reporter design included a constant 25 nt-long region at the very 5’ end of the transcript and Optimus 5 Prime may not have learned to properly model the influence of sequence elements specific to this region^42–45^. Moreover, the overall length of the tested 5’UTRs was 75 nt (25 fixed and 50 random nt) in most of our experimental assays while for mRNA technology applications it may be desirable to shorten the 5’UTR to minimize the overall transcript length and to reduce the likelihood of unintentionally including cis-regulatory information that impacts mRNA stability or translation. Second, predictive models used for mRNA therapeutics sequence design should aim to be accurate in all cell types and tissues where the therapy is expected to be functional but it is unclear whether Optimus 5 Prime predictions can generalize beyond HEK293T cells, where our reporter assays were conducted. Third, while we previously used Optimus 5 Prime to guide the design of synthetic 5’UTRs, these sequences were validated through GFP expression and ribosome loading experiments but not in a functional assay relevant to mRNA therapy or related applications^33^.

In this study, we designed *de novo* 5’UTRs for an mRNA-encoded gene editing application using Optimus 5-Prime. We first sought to characterize whether Optimus 5-Prime generalized to two new cellular contexts relevant to mRNA therapeutics. We targeted cultured hepatocellular carcinoma (HepG2) cells as a proxy for liver cells, for which protein replacement^8^, regenerative^50^, and gene editing^51^ mRNA therapies are currently being developed. Additionally, we characterized T cells, where therapeutic mRNA has been used for transient expression of chimeric antigen receptors (CAR)^52^ and to express gene editors to knock out specific receptors during manufacturing of allogeneic CAR T cells^53^. Moreover, to study 5’UTR regulation specific to the 5’ terminal region, we constructed and characterized new mRNA reporter libraries with shorter, 25nt or 50nt-long, completely randomized 5’UTRs, and used the resulting data to train a new predictor, Optimus 5-Prime(25). Finally, we designed 5’UTRs for mRNA-delivered megaTAL gene editing therapeutics. We used both versions of Optimus 5-Prime along with two design methods we recently developed: Fast SeqProp, based on gradient descent optimization^54^, and Deep Exploration Networks (DENs), based on generative neural networks^55^. We synthesized megaTAL-encoding mRNAs with our designed 5’UTRs, conducted gene editing assays in K562 cells, and found that 24 out of 29 *de novo* UTRs designed to maximize translation resulted in high editing efficiency compared to endogenous controls for two different megaTALs. Furthermore, maximum editing activity was achieved with one of the DEN-designed 5’UTRs for one of the megaTAL targets. Our results highlight the potential of current sequence design methods for mRNA therapeutics and outline limitations of our current translation predictive models.

## Results

### Optimus 5-Prime predictions generalize to cells relevant to mRNA therapeutics

We performed Massively Parallel Reporter Assays (MPRAs) to measure translation efficiency from our previously developed 5’UTR reporter library in HepG2 and T cells (**Figure 1B**), following an identical procedure as we did with HEK293T^33^ (**Methods**). Briefly, our library comprised *in vitro* transcribed (IVT) mRNAs, with a 5’UTR containing an initial constant 25nt-long segment followed by a 50nt-long fully random region, an EGFP CDS, and a 3’UTR derived from the bovine growth hormone (BGH) gene. We transfected the IVT mRNA library and, after an 8 hour incubation period, extracted cell lysates in the presence of the translational inhibitor cycloheximide, performed polysome profiling to separate polysome fractions, and sequenced each fraction. As a proxy for translation efficiency, we calculated the Mean Ribosome Load (MRL) for each 5’UTR, by multiplying the normalized read count on each fraction by the corresponding number of ribosomes^33^.

After filtering for sequences with at least 100 reads in all datasets, we obtained translation measurements from 204,803 5’UTR variants in common across five replicates over three cell types, with similar quality and read coverage (**Supplementary Figure 1A** and **B**). Analyzing a subset of the 20,000 sequences with the highest coverage, we found MRLs to be highly correlated across cell lines (**Figure 1C**, **Supplementary Figure 1C**), with coefficients of determination between cell lines (r^2^ = 0.837-0.870 for HEK293T versus T cells and r^2^ = 0.847-0.871 for HEK293T versus HepG2) comparable to those across replicates from the same cell line (r^2^ = 0.938 for HEK293T replicates and r^2^ = 0.814 for T cell replicates, **Supplementary Figure 1C**). While r^2^ decreased as we included more sequences with lower coverage, likely an artifact of decreasing data quality, their relationship across cell lines was maintained (**Supplementary Figure 1D**).

Next, we compared these measurements to Optimus 5-Prime predictions (**Figure 1D**). While the highest correlation was observed with HEK293T measurements (r^2^ = 0.937 on 20,000 sequences with the highest read coverage held out from training, **Figure 1E**), correlations with measurements in T cells and HepG2 were also high (r^2^ = 0.841 and 0.840 respectively). Prediction accuracy did not consistently increase when retraining Optimus 5-Prime individually in each cell line (**Supplementary Figure 2**) or when training a single multi-output model to predict on all cell lines simultaneously (**Supplementary Figure 3**). Finally, given that the most influential known regulatory elements are composed of three letters (AUG, CUG, GUG, etc), we investigated whether any 3-mers could have differential effects over MRL in different cell types. To this end, we trained simple 3-mer-with-position linear models on each replicate (**Supplementary Figure 4A,B**) and analyzed the resulting weights, but failed to find any cell line differences beyond those present in replicates of the same cell line (**Supplementary Figure 4C**). Together, these results show that observations from our polysome profiling MPRA in HEK293T, as well as predictions from Optimus 5-Prime, generalize to HepG2 and T cells.

*De novo* designed 5’UTRs enable high gene editing efficiency from megaTAL-encoding mRNAs Next, we used Optimus 5-Prime to design *de novo* 5’UTRs, with the goal of maximizing megaTAL expression from mRNA vectors and therefore improving gene editing efficiency. Specifically, we used megaTALs designed to disrupt two genes whose knockout in engineered T cells enhance antitumor activity^56,57^. The first megaTAL targeted exon 6 of the TGFBR2 gene, which codes for the TGF-β receptor II, a receptor for the TGF-β cytokine with prominent roles in development, regeneration, immune cell differentiation, and cancer^58,59^. Our second megaTAL targeted exon 1 of the PDCD1 gene, which codes for the signaling receptor Programmed Cell Death Protein 1 (PD-1) which acts as an inhibitory checkpoint during T cell activation^60^.

We designed 19 *de novo* 5’UTRs and selected 11 control sequences, incorporated them into megaTAL-encoding mRNAs, and quantified their performance via gene editing assays (**Figure 2A, B**). Our controls included eight sequences previously measured in our polysome profiling MPRA^33^, including four with low or medium measured MRLs as well as four selected from the top 0.02% by measured MRL (**Supplementary Figure 5A**). As additional controls, we included the 5’UTR sequences of the human VAT1 and LAMA5 genes which were identified in a prior gene editing screen as the best performing natural 5’UTRs. In previous MPRA measurements^33^ these UTRs showed high translation efficiencies similar to the commonly-used beta-globin UTR (top ∼20% among ∼17,000 short endogenous, **Supplementary Figure 5B**). We alsoincluded a minimal 5’UTR consisting of nothing more than a strong Kozak sequence^61^, which we had previously found to result in high editing efficiency. All UTRs were preceded only by the initial guanine triplet inserted by IVT.

**Figure 2.**
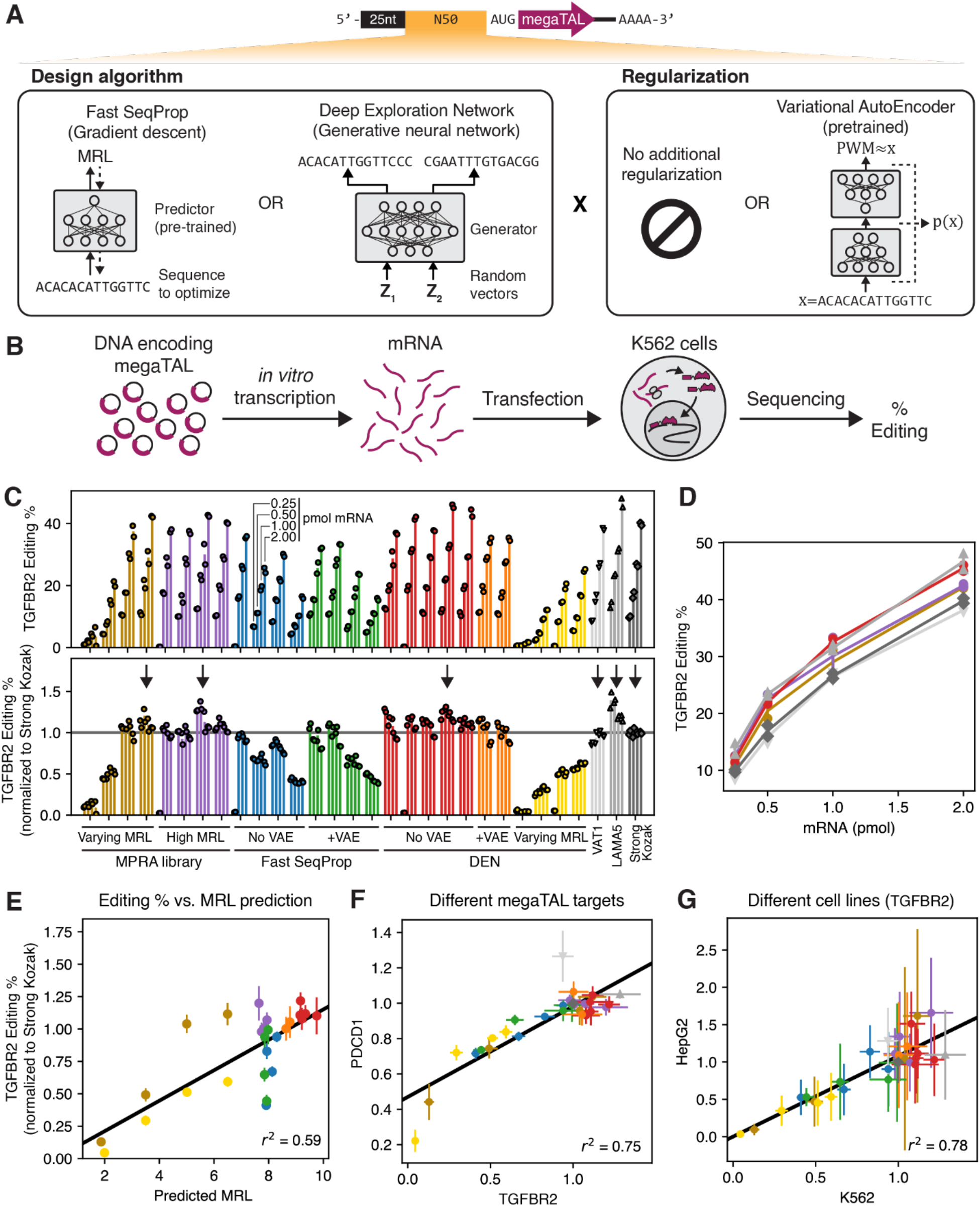
Model-based design of 5’UTRs for gene-editing mRNA therapeutics. **(A)** Top: schematic of megaTAL mRNA vector. The 5’UTR has the same architecture as in our MPRA (Figure 1B). Bottom: the variable 5’UTR region is designed via a combination of two design algorithms (Fast SeqProp^54^ or Deep Exploration Networks (DENs)^55^) and two regularization strategies (no additional regularization or Variational AutoEncoders (VAEs)^62^). **(B)** Schematic of gene editing experiment. megaTAL mRNA with each 5’UTR was individually synthesized via IVT and transfected into K562 cells. After 72 hours, DNA sequencing of the genomic region targeted by the megaTAL was sequenced and the gene editing efficiency was calculated. **(C)** Editing efficiencies for mRNAs with a megaTAL targeting the TGFBR2 gene, for 30 different 5’UTR including designs and controls. Each group of four bars represents the editing efficiency of one 5’UTR sequence transfected at four mRNA doses (0.25, 0.5, 1, or 2 pmol mRNA). Two biological replicates were performed per 5’UTR and mRNA dosage, and are represented by individual markers. Colors represent the source in the case of controls or the design method. Top: absolute editing efficiencies. Bottom: efficiencies normalized to the Strong Kozak control at the corresponding mRNA dosage. Editing efficiencies for the first High MRL MPRA control, the first No VAE Fast SeqProp design, the second No VAE DEN design, and the third Varying MRL DEN design were close to zero only at a dosage of 0.25 pmol mRNA, and were deemed to be the result of experimental error and excluded from subsequent analysis. **(D)** Absolute editing efficiency as a function of mRNA dosage for a few selected 5’UTRs, indicated with a vertical arrow in the bottom panel of **(C)**. **(E)** Kozak-normalized editing efficiency of the TGFBR2 megaTAL vs. Optimus 5-Prime predicted MRL for all designed and MPRA control 5’UTRs. **(F)** Comparison of Kozak-normalized editing efficiencies when using a megaTAL targeting the TGFBR2 or the PDCD1 genes. **(G)** Comparison of Kozak-normalized editing efficiencies for the TGFBR2 megaTAL in K562 versus HepG2 cells. In **(E)**, **(F)**, and **(G)**, each marker and error bar represent the mean and standard deviation of the Kozak-normalized editing efficiencies across all mRNA dosages for a particular 5’UTR.

*De novo* 5’UTRs were designed using either Fast SeqProp^54^ or DENs^55^ (**Figure 2A**). To preserve Optimus 5-Prime prediction accuracy, 5’UTR architecture was kept identical to the MPRA library used for training: a constant 25nt-long initial region followed by a variable 50nt segment. In Fast SeqProp, a candidate sequence is iteratively refined by following the gradient of the Optimus 5-Prime-predicted MRL with respect to a continuous representation of the sequence. By following the gradient instead of scoring multiple random mutations at each step, Fast SeqProp can design high performing sequences hundreds of times faster than simulated annealing or genetic algorithms, though it may still get stuck in local optima or overfit the predictor^54^. Although out-of-frame uAUGs are unlikely to occur in high fitness sequences, in-frame uAUGs could result in high predicted MRL but would produce an incorrect N-terminus. Therefore, we included a penalty against generated uAUGs (**Methods**). To reduce the possibility of overfitting, we scored designed sequences using an independently trained linear k-mer model (**Methods, Supplementary Figure 6**). By following this procedure, we generated ten candidate sequences, from which we randomly selected four to be tested in our gene editing assays.

To design sequences with DENs, we trained a generative neural network with the objective of maximizing Optimus 5-Prime-predicted MRL while minimizing the similarity across generated sequences (**Methods, Supplementary Figure 7A-B**). By explicitly minimizing similarity, we force the generator to capture a large section of the sequence space, thereby reducing the possibility of overfitting or getting stuck in local optima^55^. We generated 1,024 5’UTRs and selected 5 from the top 20 by predicted MRL for gene editing experiments (**Supplementary Figure 7C-E**). In addition, to validate the accuracy of the design method and predictor, we trained a DEN in “inverse regression” mode, where the generator receives an additional input that specifies a target MRL (**Supplementary Figure 8A-B**), and designed four 5’UTRs with predicted MRLs of 2, 3.5, 5, and 6.5 for experiments (**Supplementary Figure 8C-F**).

As design algorithms seek to maximize performance, they may drift into low-confidence sequence space regions of the predictor, where sequences are too dissimilar from the training data and predictions are less accurate. To prevent this, we trained a Variational Auto Encoder (VAE)^62^, a neural network that learns the marginal distribution of the training data and estimates the likelihood of any new sequence with respect to this distribution. We then used the estimated likelihood as a regularization penalty to the cost function in both Fast SeqProp and DEN (**Figure 2A, Supplementary Figure 9A-F, Methods**). Specifically, we trained a VAE on a subset of 5,000 5’UTRs selected from our MPRA dataset for their high measured MRL and read depth (**Supplementary Figure 9G**). We then designed ten additional sequences via Fast SeqProp with VAE regularization and selected four for gene editing assays. Finally, we trained a new DEN with VAE regularization, generated 1,024 sequences with high predicted MRL, and picked two from the top 10 for gene editing assays (**Supplementary Figure 10**). In summary, 19 *de novo* 5’UTRs were selected for gene editing assays, including 15 sequences designed for maximal MRL (4 with Fast SeqProp, 4 with Fast SeqProp + VAE, 5 with DEN, 2 with DEN + VAE), as well as 4 UTRs with low and medium target MRLs designed with a DEN. The sequences of all 5’UTRs tested in gene editing assays can be found in **Supplementary Table 1**.

To evaluate the performance of our designs, we synthesized IVT mRNA containing candidate 5’UTRs followed by the megaTAL CDS, transfected these at four dosage levels (0.25, 0.5, 1, and 2 pmol IVT mRNA) into K562 (lymphoblast) cells, and quantified the percentage of successful non-homologous end joining (NHEJ)-mediated gene disruption via sequencing (**Figure 2B**, **Methods**). As expected, editing efficiency increased with mRNA dosage for all 5’UTRs, with several designs exceeding 40% for the TGFBR2 and 80% for the PDCD1 megaTALs at 2 pmol mRNA (**Figure 2C** top, **Supplementary Figure 11A** top, **Figure 2D**). Editing efficiencies normalized against the Strong Kozak control (hereafter Kozak-normalized efficiencies) were highly consistent across all dosage levels (**Figure 2C** bottom, **Supplementary Figure 11A** bottom). Most of the assayed sequences showed editing efficiencies comparable to the Strong Kozak control. However, 50% of the Fast SeqProp-generated sequences showed lower editing efficiencies despite having high predicted MRL, regardless of VAE regularization. Kozak-normalized editing efficiencies were highly correlated with predicted MRL over all designed 5’UTRs (**Figure 2E**, **Supplementary Figure 11B**), although Fast SeqProp-derived sequences with low editing efficiency deviated from the linear trend the most. While we found these observations to hold for both TGFBR2 and PDCD1 megaTALs (**Figure 2F**), the specific 5’UTRs resulting in maximal editing differed: LAMA5 performed the best and VAT1 performed similarly to the Strong Kozak control with the TGFBR2 megaTAL, whereas the opposite was true for PDCD1 (**Figure 2F**). Finally, we repeated our assay in HepG2 cells and, while the general trends in Kozak-normalized efficiency were maintained (**Figure 2G**), absolute efficiency was lower and measurement variability was higher (**Supplementary Figure 12**).

### Measuring translation efficiency of short, fully variable 5’UTRs

5’UTR regulation may differ when sequence elements are placed close to the 5’ terminus. For example, various pyrimidine-rich motifs have been found to influence translation in response to stress when located within a few bases of the 5’ end^42–45^. Our previous 5’UTR MPRA was unable to interrogate this region, as a fixed 25nt segment was placed at the 5’ end to facilitate library preparation (**Figure 1B**). To overcome this limitation, and to enable design of shorter 5’UTRs, we performed polysome profiling MPRAs on two new “random-end” mRNA libraries, where the 5’UTR consisted only of a variable 25nt or 50nt region preceded only by the guanine triplet introduced by IVT (**Figure 3A**). As with our previous 50nt “fixed-end” library (**Figure 1B**), we transfected these random-end mRNA libraries into HEK293T cells and collected lysates 12h later. To compensate for a lack of a constant 5’ end for PCR-based incorporation of sequencing adapters, we used template switching (TS), wherein a reverse transcriptase derived from the Moloney murine leukemia virus appends three non-templated deoxycytosines after reaching the 5’ end of the template mRNA. Then, a template switching oligo with three riboguanines (rGrGrG) in its 3’ end binds to the non-templated overhang, thereby becoming the new reverse transcription (RT) template and providing a fixed cDNA sequence for subsequent adapter incorporation (**Figure 3A**, **Methods**). We then performed Illumina sequencing and data processing (**Methods**), and calculated MRLs from read counts as previously^33^.

**Figure 3.**
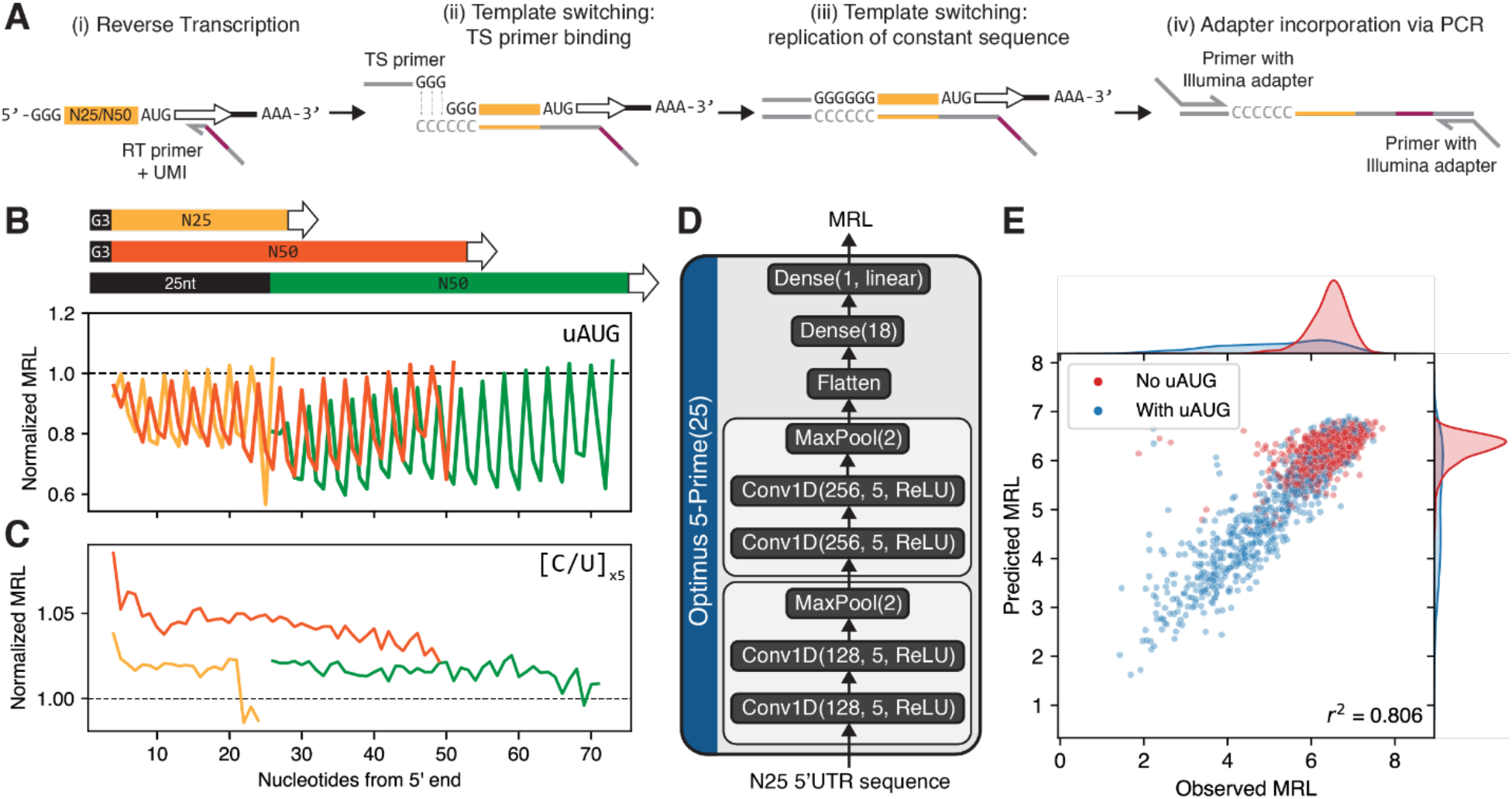
Polysome profiling MPRA on fully randomized 5’UTR libraries. New libraries contain a 25nt- or 50nt-long randomized region in the 5’UTR, preceded only by the G triplet appended by IVT. **(A)** Schematic of mRNA library preparation based on template switching (TS). (i) Reverse transcription (RT) proceeds from the EGFP CDS into the mRNA 5’end. (ii) The reverse transcriptase (RTase) adds three deoxycytosines (CCC) to the 3’ of the cDNA, to which a TS primer ending in three riboguanines (rGrGrG) binds. (iii) the RTase “switches” templates, thereby adding the reverse complement of the TS primer to the 3’ end of the cDNA. (iv) Illumina adapters are incorporated using PCR via primers that bind to the flanking constant regions. **(B-C)** Median MRL of all sequences containing a uAUG **(B)** or a 5nt-long oligopyrimidine (C or U) tract **(C)** at the indicated position from the start of the transcript, for the 25nt-(yellow) or 50nt-long (orange) randomized 5’UTR libraries, as well as our previous “fixed-end” 50nt library (green). MRL was normalized to the median of each library. **(D)** Architecture of Optimus 5-Prime(25), trained on data from the random-end 25nt MPRA library. **(E)** Performance of Optimus 5-Prime(25) on a set of 2,000 5’UTRs from the random-end 25nt library held out from training.

We performed two biological replicates with the 25nt random-end library and obtained good-quality (sum of reads across replicates greater than 100) MRL measurements from 168,000 distinct UTRs (**Supplementary Figure 13A, B**). Inter-replicate MRL correlation was good (r^2^ = 0.692 for the top 20,000 sequences by read coverage, **Supplementary Figure 13C, D**), although lower than our previous “fixed-end” 50nt library (r^2^ = 0.938, **Supplementary Figure 1D**). Similarly, we performed one replicate with the 50nt random-end library, and obtained MRL measurements from 149,000 sequences at the same quality level (**Supplementary Figure 14**). As with our previous fixed-end library, we found that 5’UTRs with uAUGs out of frame with respect to the intended AUG had significantly lower MRL compared to the median of the library and with sequences with in-frame uAUGs (**Supplementary Figure 15A**). A similar effect, although of much lower magnitude, was observed for upstream non-canonical start codons (**Supplementary Figure 15B, C**). Interestingly, MRL attenuation was noticeably lower when the uAUG was located near the 5’ end in the random-end libraries, suggesting distinct regulation at the very 5’ end compared to the rest of the 5’UTR that could not be captured in our previous fixed end library (**Supplementary Figure 15A**, **Figure 3B**).

Next, we evaluated the effect of short pyrimidine tracts (5 x C/U) at different locations within the 5’UTR on measured MRL, using data from both random-end and fixed-end library data (**Figure 3C**, **Supplementary Figure 16**). We found that pyrimidine tracts generally led to a small but statistically significant MRL increase compared to the library median (**Supplementary Figure 16**). For both libraries, we observed a noticeable decrease in effect size with increasing distance of the pyrimidine tract from the 5’ end (**Figure 3C**, **Supplementary Figure 16**). Therefore, our data is consistent with oligopyrimidine tracts at the start of the transcript resulting in slightly increased translation in HEK293 cells even in the absence of stressors.

### Predicting translation efficiency from short, fully variable 5’UTRs

We next sought to obtain a model that generates accurate predictions on 25nt-long 5’UTRs. We evaluated candidate models via their prediction accuracy on the top 2,000 sequences by read coverage from the random-end 25nt MPRA library, which showed good inter-replicate correlation (r^2^ = 0.844, **Supplementary Figure 13D**). We first tested Optimus 5-Prime, for which 25nt-long input sequences were one-hot encoded and zero padded on the left to reach the required input length. However, we found its accuracy to be relatively low (r^2^ = 0.564 for 50nt Optimus 5-Prime, **Supplementary Figure 17!**, r^2^ = 0.600 for 25-100nt Optimus 5-Prime, **Supplementary Figure 17B)**. We hypothesized that these models, trained on data from 5’UTRs with a constant 5’ region, could not properly account for differential regulatory effects that sequence elements can have when located near the start of the transcript (**Figure 3B** and **C**). Thus, we developed Optimus 5-Prime(25), a new model trained directly on the random-end 25nt MRPA data. Inspired by the convolutional network VGG-16^63^, Optimus 5-Prime(25) contains two blocks with two convolutional layers and one pooling layer each, followed by two fully connected dense layers that ultimately compute the predicted MRL (**Figure 3D**, **Methods**). This model showed good performance on the same test set which was held out from training (r^2^ = 0.806, **Figure 3E**).

### Designed short 5’UTRs enable high megaTAL-induced gene editing activity

Finally, we used Optimus 5-Prime(25) to design 14 shorter 5’UTRs for our megaTAL mRNAs. As before, we used Fast SeqProp to design ten new 5’UTRs with maximal predicted MRL, validated these against an independent k-mer linear model (**Supplementary Figure 18**, **Methods**), and randomly selected four for gene editing assays. We then trained a DEN to generate 1,024 25nt-long 5’UTRs that maximize both sequence diversity and predicted MRL, and selected 5 from the top 25 by MRL (**Supplementary Figure 19**, **Methods**). To test the effect of VAE regularization, we trained a new VAE on 5,000 high-coverage, high MRL sequences from the 25nt random-end library (**Supplementary Figure 20**, **Methods**). We then used VAE estimated likelihood as a regularization term to design ten additional 5’UTRs using Fast SeqProp (**Supplementary Figure 18**), and randomly selected two for gene editing assays. Similarly, we trained a new DEN with VAE regularization to generate 1,024 5’UTRs with maximal predicted MRL, from which we selected two from the top 10 (**Supplementary Figure 21**). As controls specific to this shorter 5’UTR design, we included four 5’UTRs from the random-end 25nt library with MRLs within the top 0.25% of the library (**Supplementary Figure 22)**.

We tested these new designs with our gene editing assay in K562 as described above (**Figure 2B**, **Methods**). As with our previous designs, editing efficiency increased with mRNA dosage (**Figure 4A**, **Supplementary Figure 23** top). When normalizing against the Strong Kozak control, we found that most of our designed 5’UTRs performed comparably to the high performing controls at all mRNA dosages, with exception of one of the Fast SeqProp designs (**Figure 4B**, **Supplementary Figure 23** bottom). Moreover, when targeting the TGFBR2 gene, one DEN-designed sequence outperformed all other UTRs at all mRNA dosages (absolute efficiency of 55.6% at 2 pmol mRNA), improving over the previous best performer LAMA5 by 18-33% (**Figure 4C**). When considering both defined-end and random-end designs, Kozak-normalized editing efficiency was highly correlated across the TGFBR2 and PDCD1 megaTALs (**Figure 4D**). However, the DEN-designed sequence that showed the highest efficiency with the TGFBR2 megaTAL performed only as well as the Strong Kozak control when combined with the PDCD1 megaTAL (absolute efficiency of 80.8% at 2 pmol mRNA), with the VAT1 control performing best in this context (absolute efficiency of 91.4% at 2 pmol mRNA, **Figure 4D**, **Supplementary Figure 23**). Finally, we repeated these experiments in HepG2 cells and found high Kozak-normalized efficiencies for all new designs, although high replication noise prevented us from identifying a single best performing sequence (**Figure 4E**, **Supplementary Figure 24**). In conclusion, by using model-based design methods with Optimus 5-Prime(25), we successfully generated *de novo* 5’UTRs that supported high gene editing activity, including one that outperformed all others for the TGFBR2 megaTAL.

**Figure 4.**
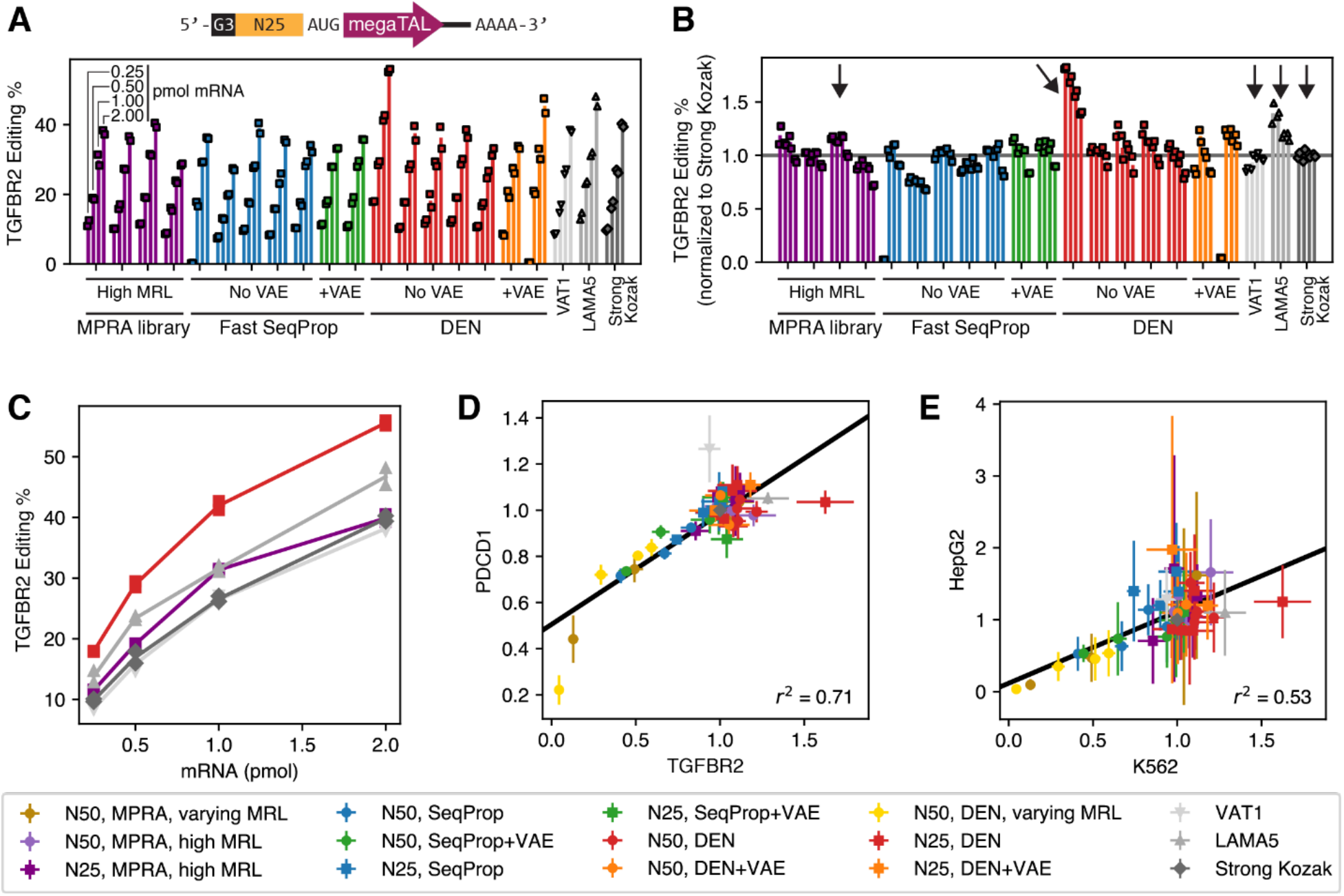
Model-based design of shorter 5’UTRs for gene editing mRNA therapeutics. **(A)** Top: schematic of mRNA vector, with a 25nt-long variable 5’UTR segment as in Figure 3. Bottom: Absolute editing efficiencies for mRNAs with a megaTAL targeting the TGFBR2 gene, for 21 different 5’UTR including designs and controls. Each group of four bars represents one 5’UTR sequence transfected at four mRNA doses (0.25, 0.5, 1, or 2 pmol mRNA). Two biological replicates were performed per 5’UTR and mRNA dosage, and are represented by individual markers. Colors represent the source in the case of controls or the design method. Editing efficiencies for the first No VAE Fast SeqProp design and the second +VAE DEN design were close to zero only at a dosage of 0.25 pmol mRNA, and were deemed to be the result of experimental error and excluded from subsequent analysis. **(B)** Editing efficiencies normalized to the Strong Kozak control at the corresponding mRNA dosage. **(C)** Absolute editing efficiency as a function of mRNA dosage for a few selected 5’UTRs indicated with a vertical arrow in **(B)**. **(D)** Comparison of Kozak-normalized editing efficiencies when using a megaTAL targeting the TGFBR2 gene vs. the PDCD1 gene. **(E)** Comparison of Kozak-normalized editing efficiencies for the TGFBR2 megaTAL in K562 cells or HepG2. In **(D)** and **(E)**, each marker and error bar represent the mean and standard deviation of the Kozak-normalized editing efficiencies across all mRNA dosages for a particular 5’UTR, and designs with the fixed-end 50nt architecture (Figure 2 and relevant Supplementary Figures) are also included.

## Discussion

In this study, we obtained MRL measurements from approximately 200,000 randomized 5’UTRs across T cells and HepG2 cells. We found that measurements were highly correlated between the two cell types and with measurements previously performed in HEK293T cells (**Figure 1C**, **Supplementary Figure 1C**). Accordingly, all measurements were accurately predicted by Optimus 5 Prime, a model trained on HEK293T cell data only. Retraining Optimus 5-Prime on data from each cell line did not consistently improve performance (**Supplementary Figure 2**). Previous MRL measurements from endogenous transcripts in a panel of cell lines^46^ and during neuronal differentiation^47^ found that 5’UTRs regulate translation mostly independently of the cellular context. Our results suggest that this is also true for synthetic 5’UTRs and in cells relevant for mRNA therapeutics.

To quantify the impact of the very 5’end of the message we then developed MPRAs with shorter, fully randomized 5’UTRs, where only three guanines at the 5’ end are kept constant due to restrictions in T7-based IVT (**Figure 3A**). This approach allowed us to observe that out-of-frame uAUGs have a smaller inhibiting effect when located close to the 5’ end (**Figure 3B**), whereas poly-C/T tracts have a small enhancing effect that increases with proximity to the 5’ end as well (**Figure 3C, Supplementary Figure 16**). Previous work has observed that multiple types of oligo pyrimidine tracts have a marked effect over translation when located at or near the 5’ end, especially in response to stress or mTOR activation^42–45^. Possibly because of these 5’-proximal regulatory effects, Optimus 5-Prime predictions were not as accurate as for the original libraries which contained a constant 25nt region at the 5’end to facilitated library preparation (**Supplementary Figure 17**). Nevertheless, we were able to train a new model, Optimus 5-Prime(25), specific to 25 nt-long fully variable 5’UTRs (**Figure 3D-E**).

Additionally, we used model-based design to generate de novo 5’UTRs for strong expression of mRNA-encoded megaTALs, supporting absolute gene editing efficiencies in K562 cells exceeding 40% when targeting the TGFBR2 gene (**Figure 2C**, **D**, **Figure 4A**, **C**) and 80% for PDCD1 (**Supplementary Figure 11**, **Supplementary Figure 23**). Notably, most of our designs resulted in high editing efficiencies, matching or exceeding the performance of 5’UTRs taken from the top 0.02% of the random MPRA libraries (**Figure 2C**, **Figure 4B**). Furthermore, one of our designs resulted in a TGFBR2 editing efficiency up to 50% higher than all controls (**Figure 4B**), though this effect was not maintained with the PDCD1-targeting megaTAL (**Figure 4D**, **Supplementary Figure 23**). Designs targeting intermediate MRLs also were found to conform to their target values, providing further support for the design approach (**Figure 2E**). Together, these results suggest that model-based design can very reliably maximize a computational predictor. Moreover, using both gene editing experiments and translation MPRAs we showed that the impact of 5’UTR sequences are broadly correlated across different cell types. but mechanisms that fully maximize expression are ORF- and possibly even cell type-dependent. Lu and coworkers recently used a high-throughput assay to test a library of 5’UTRs designed via a random forest model and a genetic algorithm^64^, and also observed that the strongest performers varied across CDSs and cell lines. However, their 5’UTRs were placed in the context of a DNA expression cassette and could therefore also influence transcription, making direct comparisons with our results difficult.

Overall, DEN-designed sequences performed better than those designed with Fast SeqProp. While the low number of sequences tested prevents us from making confident comparisons across methods, our results are consistent with Fast SeqProp being more likely to converge to solutions outside the sequence subspace where MRL predictions are accurate, because it optimizes each sequence independently. In contrast, the DEN generator network is forced to learn a representation of a large portion of the high-MRL sequence subspace, aided by the training penalty against any two generated sequences being too similar. DENs may therefore be less likely to converge to low-confidence sequence subspaces. Alternatively, these observations could also be the result of different selection strategies. With DENs, we generated 1,024 sequences per condition and chose 2-5 with the highest predicted MRL for gene editing assays. In contrast, with Fast SeqProp, we designed 10 sequences per condition but randomly chose 2-5 for megaTAL experiments.

Although designed 5’UTRs reliably mediated editing efficiencies comparable to the best controls, only one designed UTR outperformed controls by a clear margin. There are several possible reasons for this apparent upper bound on performance. First, we note that accurate out-of-distribution predictions are notoriously difficult, and the design algorithms may simply not be able to access regions of the sequence space that are not explored by the training data. Second, we can speculate that very efficient translation is a default behavior and the role of 5’UTRs is often to downregulate translation to a level that is sufficient in a specific biological context. This would be consistent with our observation that a minimal 5’UTR consisting primarily of a strong Kozak sequence results in close to maximal editing efficiencies (**Figure 2C**, **Figure 4B**). Third, it is possible that our polysome profiling assay has limited resolution at the higher end, as it is difficult to resolve polysome peaks at around 8 and higher, which may have obscured potential differences at the upper MRL end in our translation assays. Fourth, our methods do not account for 5’UTR effects on mRNA stability, which could negatively affect megaTAL expression. Previous work has showed that mRNA can be destabilized by 5’UTRs with both low^29^ and high^28^ translation efficiencies, highlighting the need for models that take into account both phenomena to optimize expression. However, our editing efficiencies were uncorrelated with mRNA half-life predictions made with Saluki, a state-of-the-art neural network predictor^35^ (**Supplementary Figure 25A and B**). Furthermore, Saluki predictions suggest that short 5’UTR regions are unlikely to affect mRNA half-life dramatically (**Supplementary Figure 25C**). However, since Saluki was trained on endogenous transcript datasets, its predictive accuracy on synthetic sequences is currently unknown. Finally, processes downstream of megaTAL translation, including nuclear import and DNA cleavage kinetics, may limit gene editing efficiency.

In summary, we used deep learning methods to engineer 5’UTRs that resulted in strong gene expression of a genome editing mRNA therapy. While most of our designs enabled strong gene editing activity, our ability to identify and design outperformers for all ORFs and cell lines of interest is still restricted. This limitation may be overcome by training predictive models on data from MPRAs that interrogate the interactions between 5UTR, ORF, and 3’UTR sequences, reliably measure top-end translation efficiency, and potentially characterize other biomolecular phenomena such as mRNA stability.

## Methods

### Polysome profiling in HepG2 cells

HepG2 cells were cultured in EMEM + 10% FBS. The cells were in culture prior to the experiment, passage conditions were 20 mL media and cells at 2e5/mL into a T-75 flask, and cells were allowed to expand for 3 days prior to transfection. Cells used were at passage 6. IVT mRNA corresponding to the 5’UTR 50nt “fixed-end” library was taken from our previous work^33^. 1 ug IVT mRNA was transfected into one million cells with the Lonza 4D Nucleofector, following the manufacturer’s protocol. Cell lysis was performed 8 hours later, followed by polysome profiling, library preparation, and Illumina sequencing as previously described^33^.

### Polysome profiling in T cells

T cells were enriched from peripheral blood mononuclear cells (PBMCs) isolated via Ficoll-Paque gradient centrifugation from healthy donors. Activation was performed with anti CD3/CD28 antibodies in the presence of IL-2. Salt solution and lysis buffer for polysome profiling were prepared as previously described^33^. IVT mRNA corresponding to the 5’UTR 50nt “fixed-end” library synthesized as part of our previous work^33^ was used here. 1 ug IVT mRNA was transfected into one million cells with the Lonza 4D Nucleofector, following the manufacturer’s protocol. Transfected cells were plated in T cell growth medium (TCGM) at 1 million cells/mL and incubated at 37C, 5% CO_2_. 8 hours later, cycloheximide was added dropwise to the media to a final concentration of 100 ug/mL and incubated for 5 additional minutes. Cells were then spined down at 1500 rpm for 5 min and the supernatant was discarded. 300 uL of cold lysis buffer was used to resuspend the cells, and the mixture was incubated on ice for 10 minutes. Cells were then triturated by passing the mixture through a 25-gauge needle ten times. The mixture was spined down at 16,000 rpm for 5 min at 4C and the supernatant was transferred to another tube. 1.5 uL of 1U/uL DNAse was added, the mixture was set on ice for 30 minutes, and stored at -80C. Polysome profiling, library preparation, and Illumina sequencing were performed as previously described^33^.

### Construction of random-end 5’UTR MPRA libraries

Our previously constructed vector pET28-IVT-Fixed-AgeI-EGFP-NheI^33^, containing a T7 promoter followed by the 25nt-long defined 5’UTR prefix and the EGFP CDS, was amplified with primers “Bri035 FP EGFP START” (TGGGCGAATTAAGTAAGGGC) and “Bri042_RP_T7” (CCCTATAGTGAGTCGTATTAATTTCGCG). This resulted in a linear dsDNA backbone with the complete original vector sequence except for the 25nt-long defined 5’UTR fragment. Random 50nt- and 25nt-long 5’UTRs were introduced by assembling 200 ng of backbone with 10 pmol of primer Bri036_T7_N50_ATG (TAATACGACTCACTATAGGGNNNNNNNNNNNNNNNNNNNNNNNNNNNNNNNNNNNNNNNNNNNNNNNNNNATGGGCGAATTAAGTAAGGG) or Bri043_T7_N25_ATG (GCGAAATTAATACGACTCACTATAGGGNNNNNNNNNNNNNNNNNNNNNNNNNATGGGCGAATTAAGTAAGGGCGAGGAGCTGT), respectively, using the NEBuilder HiFi DNA Assembly Master Mix (NEB). 20 uL reactions were incubated at 50C for 1 hour to increase assembly yield. Next, water was added to reach a total volume of 100 uL per reaction, and purification was performed using the column-based DNA Clean & Concentrator 5 (Zymo) with elution in 7uL dH2O. Each library was split in two and transformed separately into 5-alpha cells (NEB, 3.5uL DNA in 35uL cells). Plasmid library isolation, IVT template synthesis, and IVT were performed as previously described^33^.

### Polysome profiling of random-end 5’UTR libraries

Transfection, cell lysis, polysome profiling, and RNA extraction were performed as previously described^33^. Extracted RNA was eluted in 11 uL RNase-free water. Library preparation comprised reverse transcription, template switching, and qPCR amplification, all of which were performed separately for each polysome fraction. For reverse transcription, we first mixed 10.5 uL of purified RNA with 2 uL of 2 uM RT primer (AGGGACATCGTAGAGAGTCGTACTTANNNNNNNNNNAGATGAACTTCAGGGTCAGC, where Ns comprise the UMI) and 2uL of 10mM dNTP mixture (NEB), incubated the mixture at 65C for 5 minutes, and placed on ice for at least one minute. Then, we added 4 uL Maxima RT buffer, 0.5 uL Superase-In RNAse inhibitor (Thermo Fisher), and 1 uL Maxima RT RNaseH minus Enzyme (Thermo Fisher), incubated at 50C for 15 min, then 85C for 10 min, and transferred back to ice. Next, we added 1 uL RNase (from bovine pancreas, DNase free, Roche) and 1 uL RNase H (NEB) and incubated at 37C for 15 minutes. Finally, the product was cleaned with KAPA beads (Roche) with a 3x ratio of beads to DNA volume, and resuspended in 25 uL dH2O. For template switching, we added 8 uL Maxima RT buffer, 6 uL 50% PEG-8000, 0.5 uL Superase-In, 1 uL Maxima RT RNaseH minus enzyme, and 0.5 uL of 10 uM template switching oligo (AAGCAGTGGTATCAACGCAGAGTACATrGrGrG). This mixture was incubated at 42C for 30 minutes, then 85C for 10 minutes. Next, 1uL RNase (Roche) and 1uL RNase H (NEB) were added and the mixture was incubated at 37C for 15 minutes. Finally, the product was cleaned with 3x KAPA beads and resuspended in 20 uL dH2O. qPCR amplification was performed using the KAPA HiFi master mix (Roche) with forward primers AATGATACGGCGACCACCGAGATCTACAC[8nt-long index barcode]AGCGTGACAGGGACATCGTAGAGAGTCGTACTTA and reverse primers CAAGCAGAAGACGGCATACGAGAT[8nt-long index barcode]AAGCAGTGGTATCAACGCAGA, where the barcodes were specific to each polysome fraction. qPCR reactions were stopped before reaching saturation and purified via gel extractions. Prepared and barcoded libraries corresponding to all polysome fractions of both random-end 25nt replicates and the one 50nt replicate were pooled into a single library for sequencing. Sequencing was performed in an Illumina NextSeq 500 with the NextSeq 500/550 v2 High Output 75 cycle kit. The following custom primers were used: read 1: GCTCCTCGCCCTTACTTAATTCGCCCAT, read 2: CACCTACGGCAAGCTGACCCTGAAGTTCATCT, index 1: CCCATGTACTCTGCGTTGATACCACTGCTT, index 2: TAAGTACGACTCTCTACGATGTCCCTGTCACGCT. Number of cycles were as follows: read 1: 59, read 2: 10, index 1: 8, index 2: 8. Read 1 data should contain the reverse complement of the variable 5’UTR followed by the reverse complement of the template switching oligo, whereas read 2 should contain the random UMI.

### Processing of polysome profiling sequencing data

Sequencing data from the “fixed-end” MPRAs in HepG2 cells and T cells (**Figure 1**) was processed identically to our previous HEK293T data^33^. For the “random-end” MPRAs (**Figure 3**), we first generated fastq files from the raw instrument output via bcl2fastq with the following options: --no-lane-splitting --minimum-trimmed-read-length 9 --mask-short-adapter-reads 9 --ignore-missing-bcls. We then used a custom python script to retain reads with a mean q-score greater than 25 and where the 3’ end of the read 1 sequence matched the expected template switching oligo with a maximum edit distance of 5. We then used starcode-umi to collapse UMIs. Finally, we calculated MRL from UMI counts on all polysome fractions as described before^33^.

### Synthesis of megaTAL mRNA

Briefly, ultramers were synthesized encoding the T7 sequence, a 5’ UTR, and the first 20 bases of the megaTAL CDS. Template for in vitro transcription was generated via PCR with the 5’ UTR-containing ultramers as the forward primer and an ultramer encoding the last 20 bases of the megaTAL CDS and a 125-base polyA tail as the reverse primer. Following plasmid degradation via DpnI, the resulting amplicon was isolated and purified using Ampure beads (Beckman Coulter). In vitro transcription was performed with ARCA co-transcriptional capping. Following DNase treatment to remove residual template, the resulting mRNA was purified using RNase-free Ampure beads. mRNA was run on the Fragment Analyzer (Agilent) to verify expected size and purity, then normalized to 500 nM and stored at -80 C until needed.

### MegaTAL gene editing assay

mRNA at amounts ranging from 2 pmol to 0.25 pmol was electroporated in duplicate into 100,000 K562 or HepG2 cells per well using the Lonza 4D Nucleofector 96-well shuttle attachment. Electroporation conditions were optimized for mRNA transfection of the respective cell type. Following electroporation, cells were cultured at 37C for 72 hours. Cells were lysed in Viagen DirectPCR lysis reagent (cell) following manufacturer protocol to extract gDNA.

### Assessment of gene editing efficiency by amplicon sequencing

Amplification of a ∼150 bp region surrounding the megaTAL target site was performed in two PCR reactions. In PCR1, 1.5 uL of genomic DNA was amplified in 30 cycles using gene-specific primers containing Illumina overhangs. In PCR2, P5/P7 sequences and unique combinations of i5 and i7 index sequences were appended to yield dual-indexed amplicons in 10 PCR cycles. Samples were pooled, cleaned up with ampure xp beads, and normalized to 16 pM, then run on an Illumina MiSeq with 25% PhiX. BCL data was converted to fastq format and paired ends were merged with PEAR. Reads were demultiplexed and aligned using bowtie2. Editing frequency was calculated as the number of reads that contained insertions or deletions that included part of the 10 base window around the expected breakpoint divided by the total reads with a MAPQ score >20 and quality score >30.

### Training of Optimus 5-Prime (25)

For every 5’UTR sequence in the random-end 25nt library, a weighted averaged MRL was obtained across replicates, with weights given by the total number of UMI reads per replicate. Sequences were then sorted by read depth, and those with fewer than 100 reads were discarded. The top 2,000 sequences by read depth were held out for testing, the next 2,000 were used for validation/early stopping, and the remaining 193,341 were used for training.

Model training and evaluation were done in Python 3 with tensorflow 2. Optimus 5-Prime(25) architecture was based on VGG-16^63^: it contains a number of convolutional blocks – each with two convolutional layers, one max pooling layer with size and stride 2, and one dropout layer – followed by a fully connected dense layer and a final linear node that computes the MRL. All activations except for the final node are ReLU. The number of convolutional filters in each block is twice the number of filters in the previous block. Models were trained using an MSE loss, and early stopping based on the validation loss was used. Hyperparameter tuning was performed with Amazon Sagemaker using their default Bayesian strategy, with the following parameter ranges: number of convolutional blocks: 1 - 5, kernel size: 2 - 7, number of filters in the first convolutional block: 16 - 128, convolutional dropout: 0 - 0.5, number of units in the final dense layer: 10 - 100, dropout: 0 - 0.5. The final architecture is shown in **Figure 3D**.

### 5’UTR sequence design using Fast SeqProp

All relevant code was run in Python 3 with keras 2.2 and tensorflow 1.15. Sequence design with Fast SeqProp was done as in our previous publication^54^. The following is a summary of design details specific to this study.

For Fast SeqProp designs with the 50nt “fixed-end” architecture (**Figure 2**), we used a “retrained” version of Optimus 5-Prime initially trained on the fixed 50nt MPRA data and finetuned on sequences designed to maximize MRL that contained long poly-U stretches and ultimately underperformed^33^. The loss function to minimize was the sum of a fitness loss plus a sequence loss. The fitness loss was set to the negative of the predicted MRL. The sequence loss was set to the number of occurrences of AUGs across the generated sequence. We also included a term that was set to one if an “UG” was present at the beginning of the designed region and zero otherwise. This is to penalize an initial uAUG since the last nucleotide of the fixed-end region was A. When VAE regularization was used, an additional term corresponding to the VAE loss was computed by passing the VAE-estimated p_VAE_(seq) through a margin function max(0, margin_vae – log(p_VAE_(seq))), where margin_vae = -30. The VAE loss was multiplied by a weight of 0.2 and added to the overall loss function. The number of gradient updates (iterations) was 20,000 for the non-VAE designs and 5,000 for the VAE-regularized designs.

Fast SeqProp designs with the 25nt variable-end architecture (**Figure 4**) were done as above, with the following changes: 1) we used Optimus 5-Prime(25) (**Figure 3**), 2) we did not include a penalty for an initial UG dinucleotide, 3) margin_vae was set to -15.6, and 4) the vae loss weight was set to 0.4.

### Training of k-mer models for validation of Fast SeqProp designs

A linear k-mer model was trained on a subset of the HEK293T 50nt fixed-end MPRA data and used as an additional oracle to “validate” Fast SeqProp designs (**Supplementary Figure 6**). Models were trained using the linear_model module of the scikit-learn python package. As training data, MPRA sequences were filtered by discarding those with fewer than 250 reads, those with uAUGs, and those starting with UG to avoid creating an uAUG with the last nucleotide of the fixed 5’ region which was an A. From the remaining 125,931, sequences, 100,000 were used for training and 25,931 for testing. Next, counts of 2, 3, 4, 5, and 6-mers were calculated, and log2(1 + kmer counts) were computed, resulting in a feature vector of size 5,456. A Lasso model with α = 0.001 was trained on a random subset of 50,000 sequences from the training data, resulting in 272 non-zero feature weights. Finally, a Ridge regression model with α = 0.0 was trained using only the non-zero features from the Lasso regression model. Performance on the test set is shown in **Supplementary Figure 6B** (Pearson r = 0.5213), and model predictions for sequences designed with Fast SeqProp are shown in **Supplementary Figure 6C.**

A similar model was trained for validating Fast SeqProp designs with the 25nt random-end architecture (**Supplementary Figure 18**). Training sequences were taken from the 25nt random-end MPRA data, then retained only if their read count was greater than 150 and if they did not contain uAUGs. From the remaining 81,552 sequences, 70,000 were used training and 11,552 for testing. Calculation of feature vectors from kmer counts, Lasso regression, and Ridge regression were performed as above. 268 features with nonzero weights resulting from Lasso regression were used for Ridge regression. Performance on the test set is shown in **Supplementary Figure 18A** (Pearson r = 0.4094), and model predictions for sequences designed with Fast SeqProp are shown in **Supplementary Figure 18B**.

### 5’UTR sequence design using Deep Exploration Networks

All relevant code was run in Python 3 with keras 2.2 and tensorflow 1.15. Sequence design with Deep Exploration Networks (DENs) was performed as in our previous publication^55^. In total, we trained five DENs: two for maximizing MRLs in a fixed-end 50nt architecture (**Figure 2**), without (**Supplementary Figure 7**) and with (**Supplementary Figure 10**) VAE regularization, one for designing sequences with submaximal MRLs (“inverse regression”, **Supplementary Figure 8**), and two for maximizing MRLs in a variable-end 25nt architecture (**Figure 4**), without (**Supplementary Figure 19**) and with (**Supplementary Figure 21**) VAE regularization. DEN generator architectures, a short description of the loss function components, and training parameters can be found in the corresponding supplementary figures.

### Training of Variational AutoEncoders

VAE training and evaluation were done in Python 3 with keras 2.2 and tensorflow 1.15. Model architectures for the fixed-end 50nt and the random-end 25nt VAEs are shown in **Supplementary Figure 9** and **Supplementary Figure 20** respectively. A high level description of how VAEs are trained and evaluated can be found in **Supplementary Figure 9A** and **B**. Detailed information on the loss function, including equations and rationale for each term, can be found in our previous publication^55^. For the fixed-end 50nt VAE, sequences for training and testing were extracted from our published fixed-end 50nt MPRA dataset^33^, by first filtering by read coverage (> 2,000 reads) and then randomly selecting 5,000 (train) and 1,000 (test) sequences from the top 10,000 by MRL (∼top 25%). For the random-end 25nt VAE, sequences for training and testing were extracted from the random-end 25nt MPRA dataset (**Figure 3**), by filtering by read coverage (> 500 reads) and then randomly selecting 5,000 (train) and 1,000 (test) sequences from the top 10,000 by MRL (∼top 25%).

### Code Availability

Models and code are available at https://github.com/castillohair/paper-5utr-design.

---

**Supplementary Figure 1.**
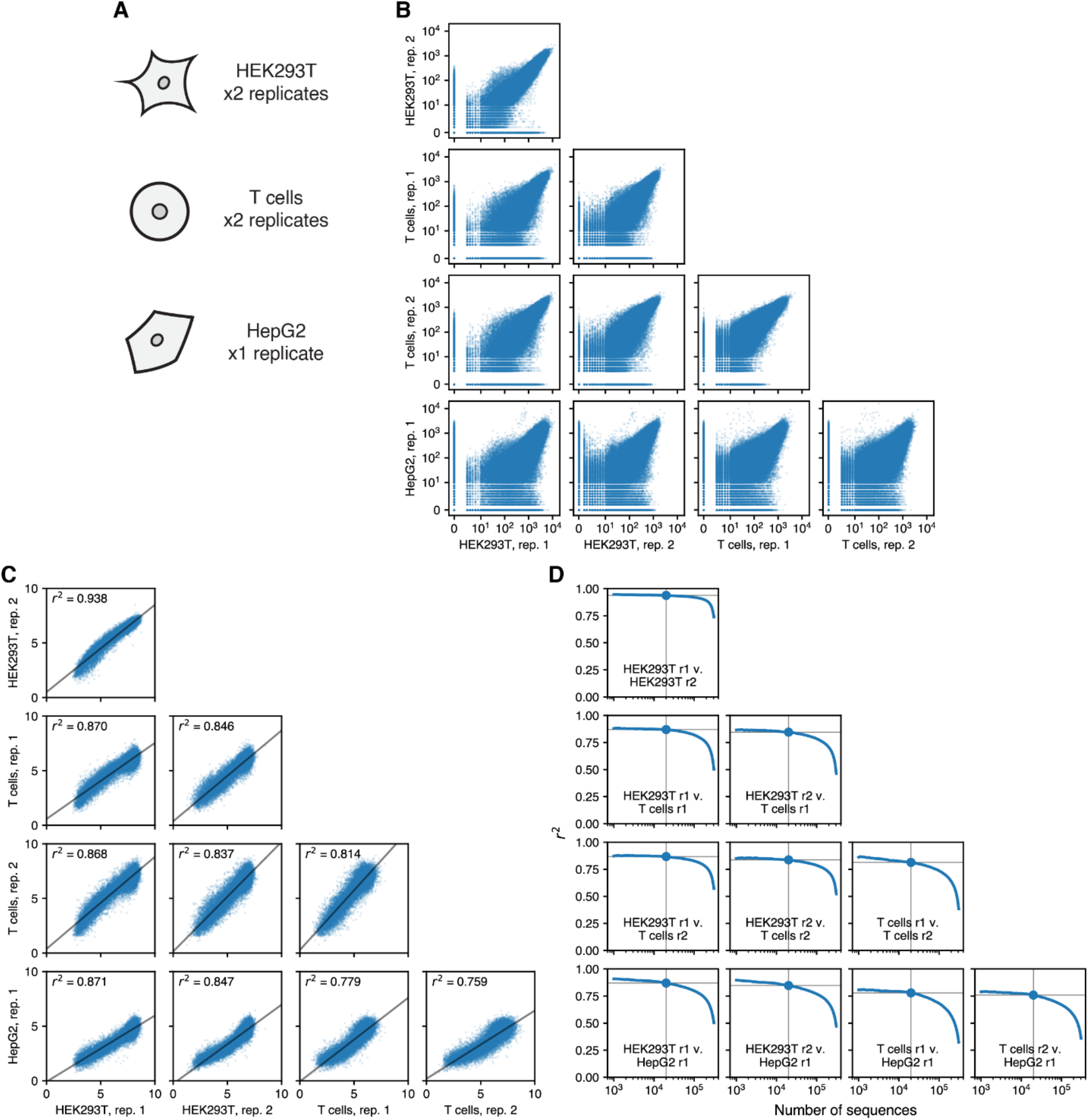
Comparison of polysome profiling MPRA data in HEK293, T cells, and HepG2. **(A)** Cell lines and number of biological replicates. **(B)** Comparison of number of reads for each 5’UTR sequence across all pairs of replicates. **(C)** MRL comparison for the 20,000 5’UTR sequences with the highest minimum read coverage across all replicates. A regression line for each pair is shown in black along with the coefficient of determination r^2^. **(D)** r^2^ as a function of the number of sequences used. Sequences were sorted by the minimum number of reads across all replicates in descending order. Then, the top x sequences (x axis) were used to calculate a corresponding r^2^ value (y axis). The large marker and the gray lines indicate the number of sequences and r^2^ in **(C)**.

**Supplementary Figure 2.**
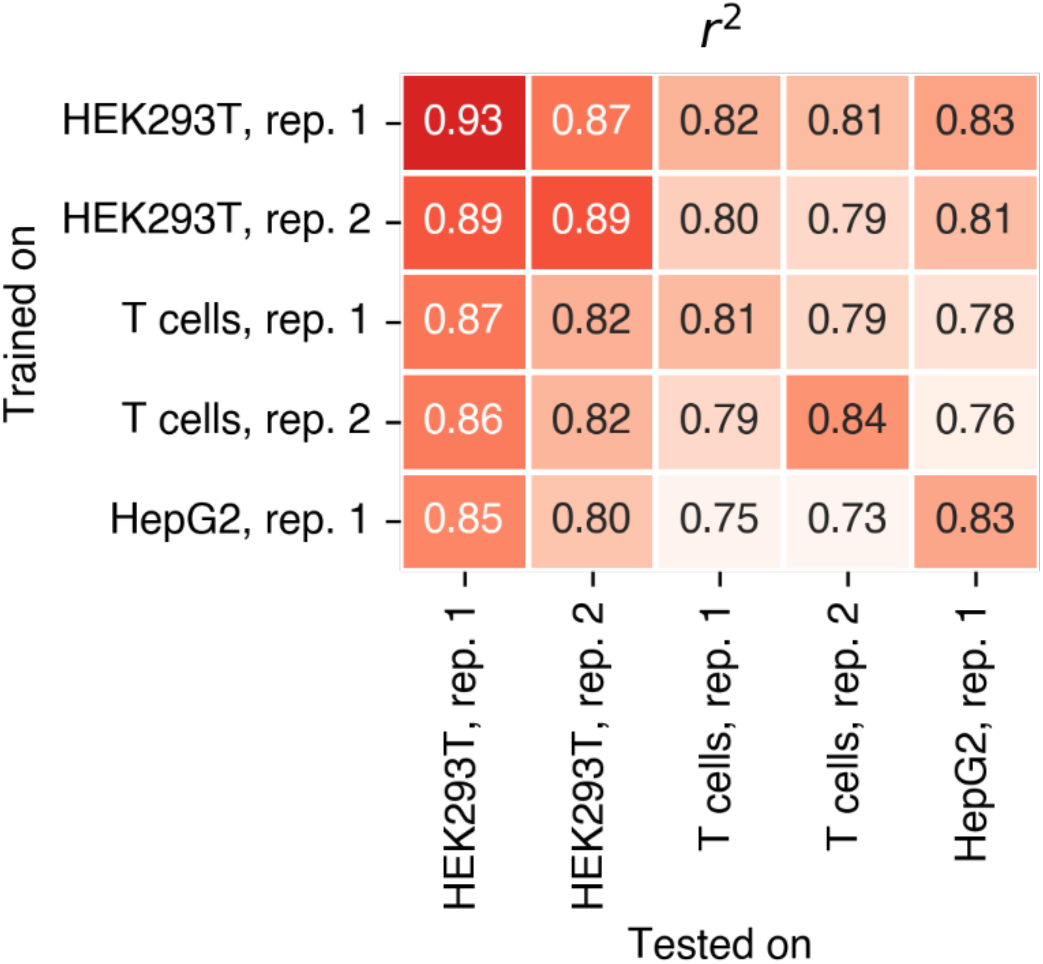
Optimus 5-Prime performance when retrained on cell type-specific polysome profiling data. The top 20,000 5’UTR sequences with the highest minimum read coverage across all cell type replicates were separated for testing, and the remaining sequences were used for training. For every cell line and replicate indicated in each row, Optimus 5-Prime was retrained from scratch on the training dataset after filtering for sequences with more than 200 reads. Then, MRL predictions on the test dataset were generated and compared with measurements from each cell line and replicate in each column.

**Supplementary Figure 3.**
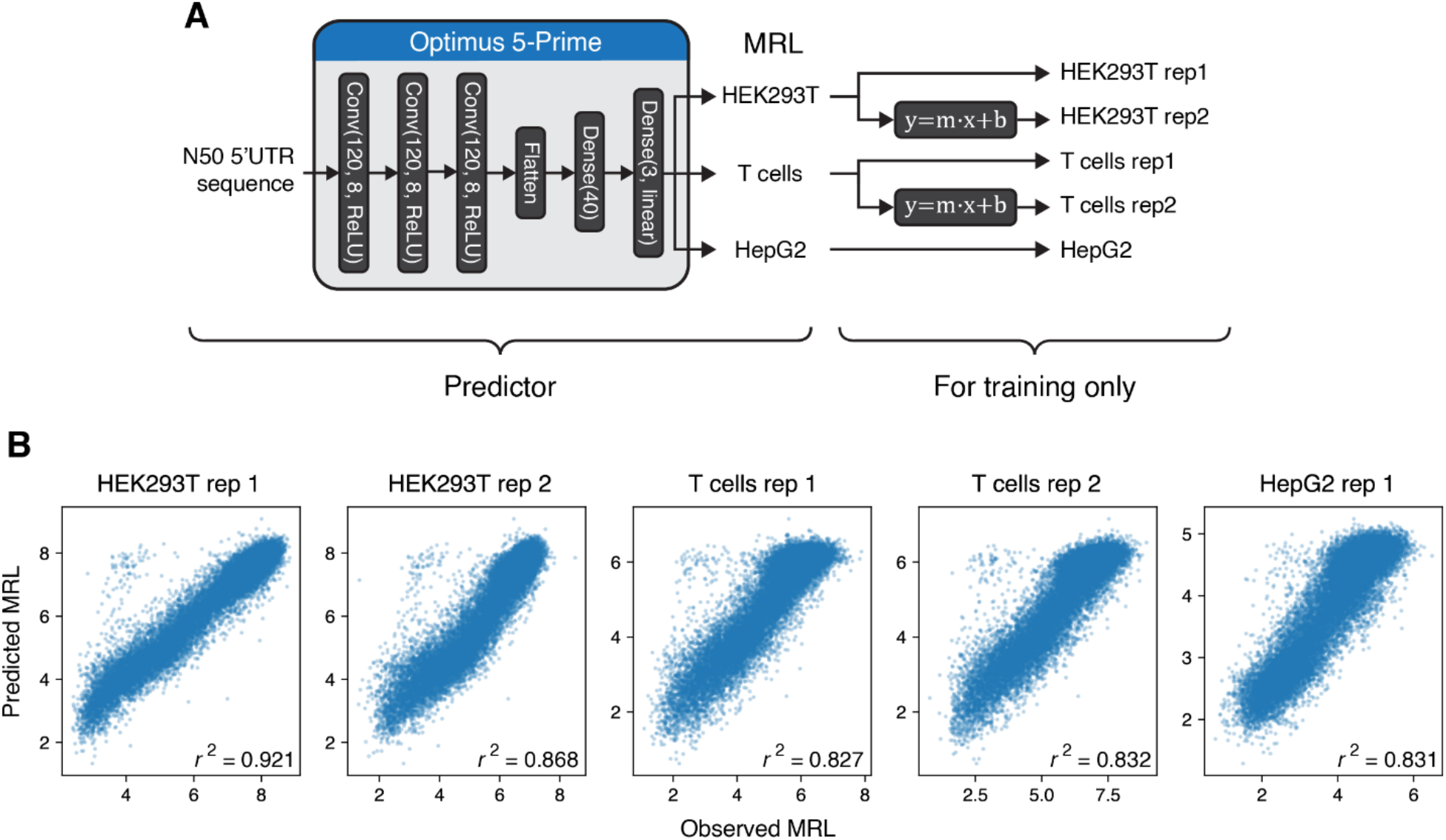
Performance of a multi-output version of Optimus 5-Prime. **(A)** Model diagram. The architecture is the same as the original Optimus 5-Prime, but with a final dense layer with three outputs corresponding to each cell type. During training, we added an additional final layer containing learnable linear scalings to account for systematic bias of replicates of the same cell type. This final layer is not saved with the model after training. **(B)** Model performance when compared to MRL measurements in all cell type replicates, on a held-out test dataset containing 20,000 sequences with the highest minimum read coverage across all replicates.

**Supplementary Figure 4.**
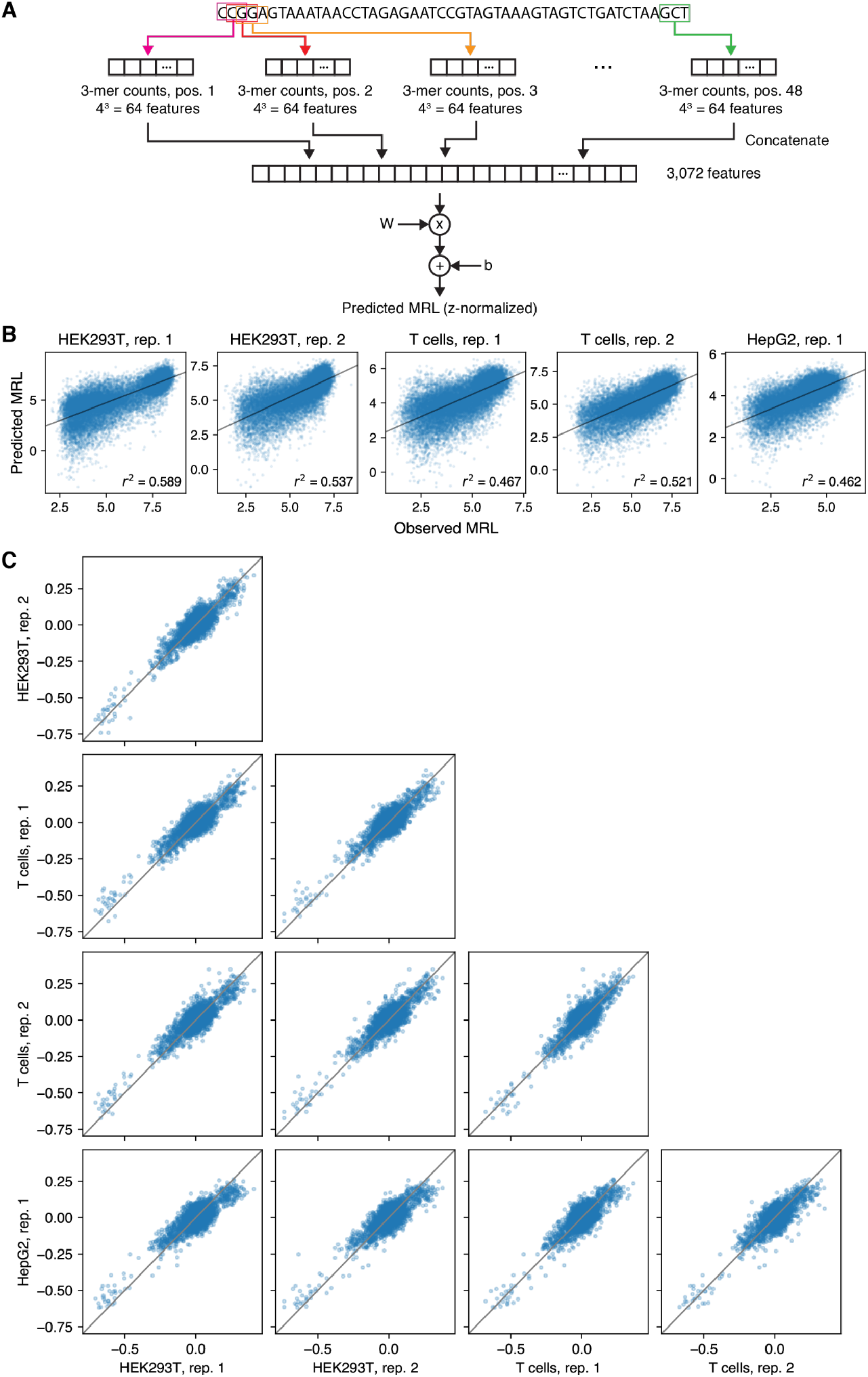
Analysis of 3-mer with position models across all cell type replicates. **(A)** Model schematic. Five models, one per cell line and replicate, were trained on z-normalized data using Ridge regression and a regularization coefficient of 1e-5. Regression weights and bias are represented by the 3,072-long vector W and the scalar b. **(B)** Model performance. Models were evaluated on the 20,000 sequences with highest read coverage on each cell line and replicate, which were held out from training. **(C)** Comparison of model parameters (3,072 weights + bias) across all five models.

**Supplementary Figure 5.**
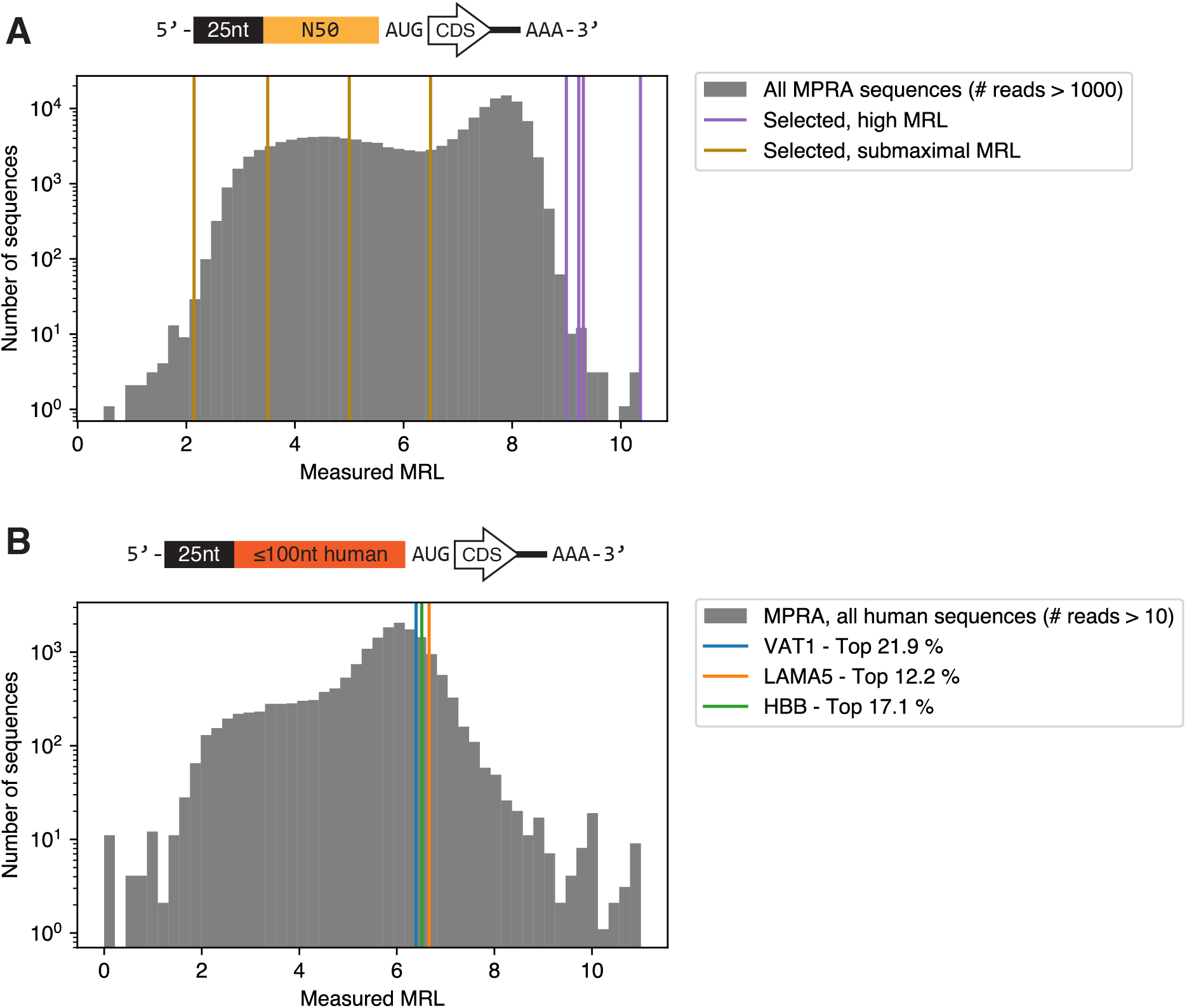
Controls for the megaTAL gene editing assays. **(A)** Selection of controls from the fixed-end MPRA library. For the “high MRL” control set, starting from the HEK293T MPRA library we excluded sequences with a read count lower than 1,000 or if they contained uATGs or started with TG. The remaining sequences were sorted by MRL, and four from the top twenty were selected. For the “submaximal MRL” control set, we similarly filtered by read count and excluded sequences with TG at the start, and selected four sequences with MRLs close to 2, 3.5, 5, and 6.5. Top: architecture of the MPRA library. Bottom: Histogram of a high-coverage (# reads > 1,000) subset of the MPRA library, along with the MRLs of all eight selected sequences. **(B)** Selection of controls from a library of short (<=100bp) human 5’UTRs measured in our previous polysome profiling study^33^. Top: architecture of the MPRA library. Bottom: histogram of a subset (16,779 sequences with # reads > 10) of the human 5’UTR library, along with the MRLs of the two selected sequences (VAT1, LAMA5) and the hemoglobin beta (HBB) 5’UTR commonly used in mRNA therapeutics. The legend indicates the MRL percentile of these three 5’UTRs compared to the rest of the library. In the megaTAL experiments, human 5’UTR controls did not include the initial constant 25nt, but had a consensus Kozak sequence (GCCACC) appended at their 3’ end. See TABLE for full sequences.

**Supplementary Figure 6.**
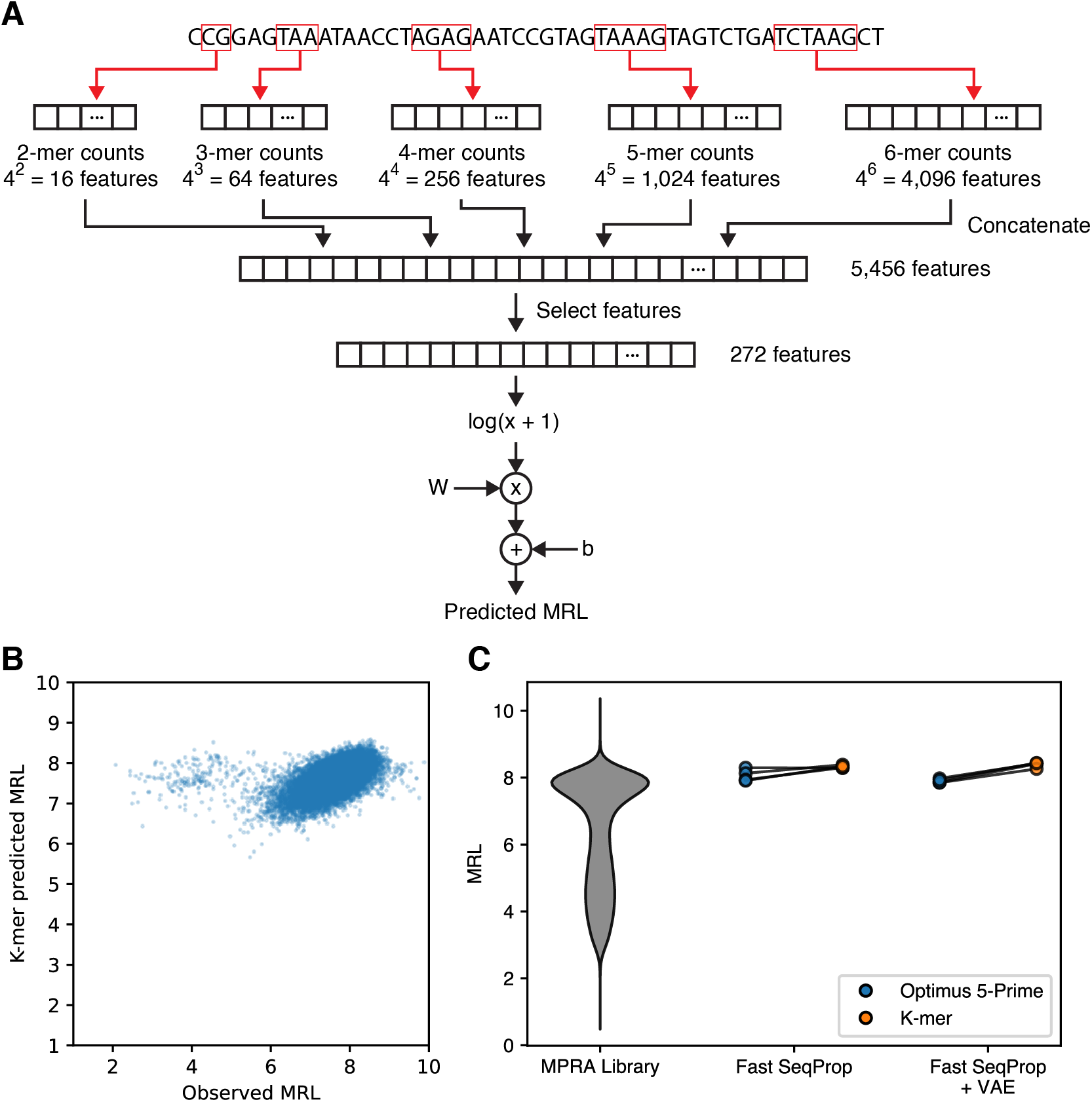
Using a linear k-mer model to validate fixed-end 50nt-long Fast SeqProp designs. **(A)** k-mer model architecture. W is a 272-long weight vector, and b is a scalar bias. See **Methods** for a description of the model and training procedure. **(B)** Observed vs. predicted MRL on a held-out set of 25,931 5’UTRs with no uAUG and greater than 250 reads. Pearson r = 0.5213 **(C)** Comparison of predicted MRL for the four sequences designed via Fast SeqProp and the four sequences designed via Fast SeqProp with VAE regularization, when using Optimus 5-Prime or the k-mer model from panel **(A)**. A violin plot of the entire MPRA library is shown on the left for comparison. K-mer model predictions from the designed sequences are within the top 1% compared to equivalent predictions on the entire test set shown in **(B)**.

**Supplementary Figure 7.**
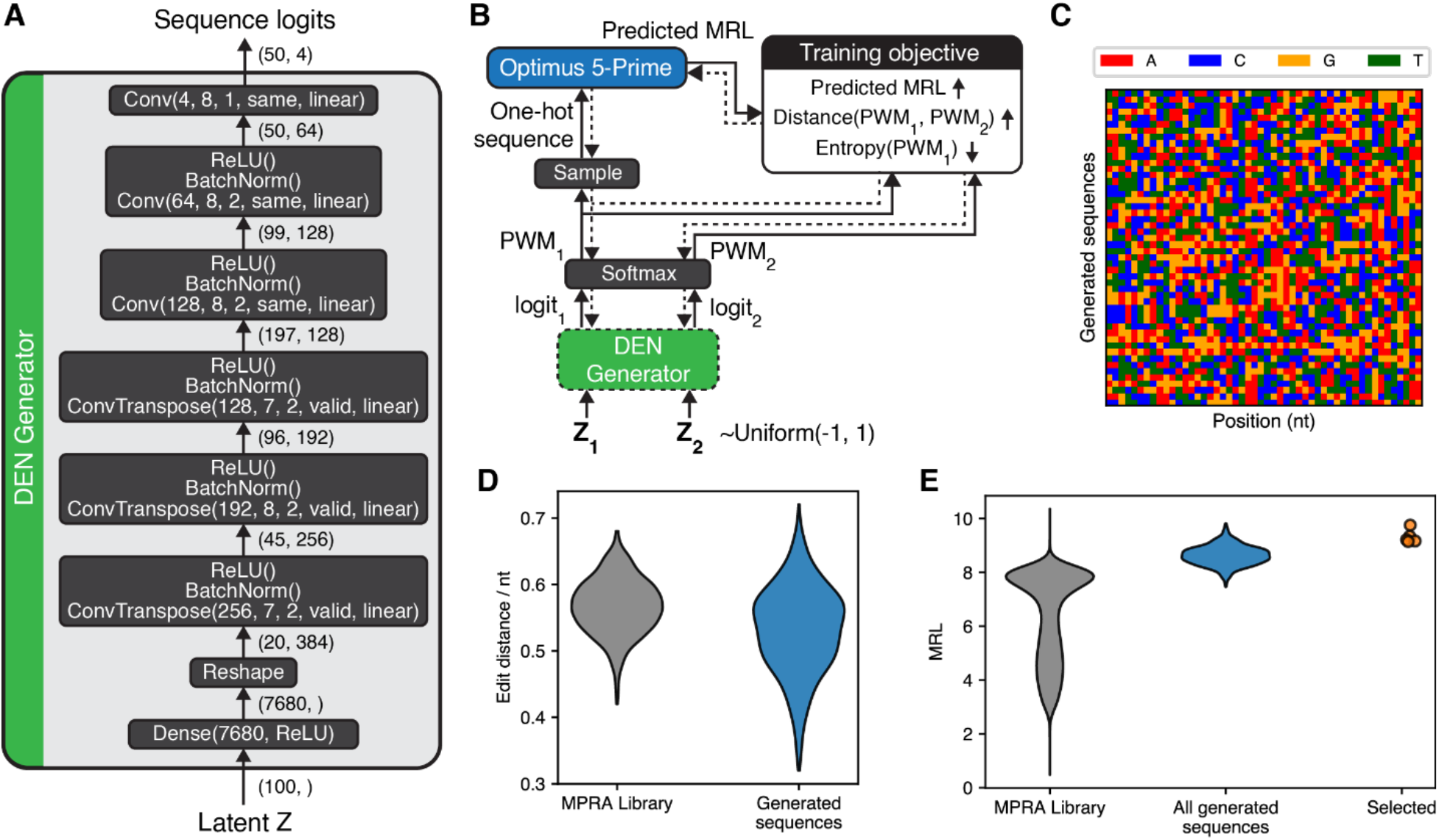
50nt 5’UTR design using Optimus 5-Prime and a Deep Exploration Network (DEN). **(A)** Architecture of the DEN generator network, which takes a continuous-valued 100-dimensional latent vector and returns a 50×4-dimesional continuous-valued logit representing a sequence. Convolutional and Transpose convolutional layers are represented as *Conv(F,W,S,P,A)* and *ConvTranspose(F,W,S,P,A)*, where *F* and *W* are the number and size of convolutional filters, *S* is the stride, *P* is the padding, and *A* is the activation function. **(B)** DEN training schematic. Only the DEN generator’s weights are optimized via gradient descent. As the predictor, we used a “retrained” version of Optimus 5-Prime, initially trained on the fixed 50nt MPRA data and finetuned on sequences we previously designed to maximize MRL that ultimately underperformed^33^. At any iteration, two sequence logits are generated from two uniformly random seed vectors. Optimus 5-Prime is used to obtain an MRL prediction from a one hot-encoded sequence sampled from *logit*_1_. The training objective to minimize is the weighted sum of the following components: 1) a fitness component set to −*MRL*, 2) a similarity component calculated from both PWMs as follows *max*(0, *CosineSimilarity*(*PWM*_1_, *PWM*_2_)*margin_similarity_*), and 3) an entropy component set as the Shannon Entropy of *PWM*_1_ up to a margin *max*(0, *margin_entropy_* – (2 – *entropy*)) to ensure*PWM*_1_ is close to a one hot-encoded sequence. Weights for the fitness, similarity, and entropy components of the loss function were 0.1, 5, and 1. Margin values for the similarity and entropy terms were 0.3 and 1.8. Training was performed for 250 epochs. **(C)** Colormap representation of 50 5’UTR sequences randomly chosen from the 1,024 generated by the trained DEN. Each row represents a separate sequence, and color indicates nucleotide identity. **(D)** Distribution of edit distances per nucleotide, for 500 random pairs chosen from the MPRA library (left) or the 1,024 sequences generated by the DEN. **(E)** Distribution of MRLs measured from the MPRA library (left), or predicted by Optimus 5-Prime on all 1,024 DEN-generated sequences (middle) and the four selected sequences from this set (right).

**Supplementary Figure 8.**
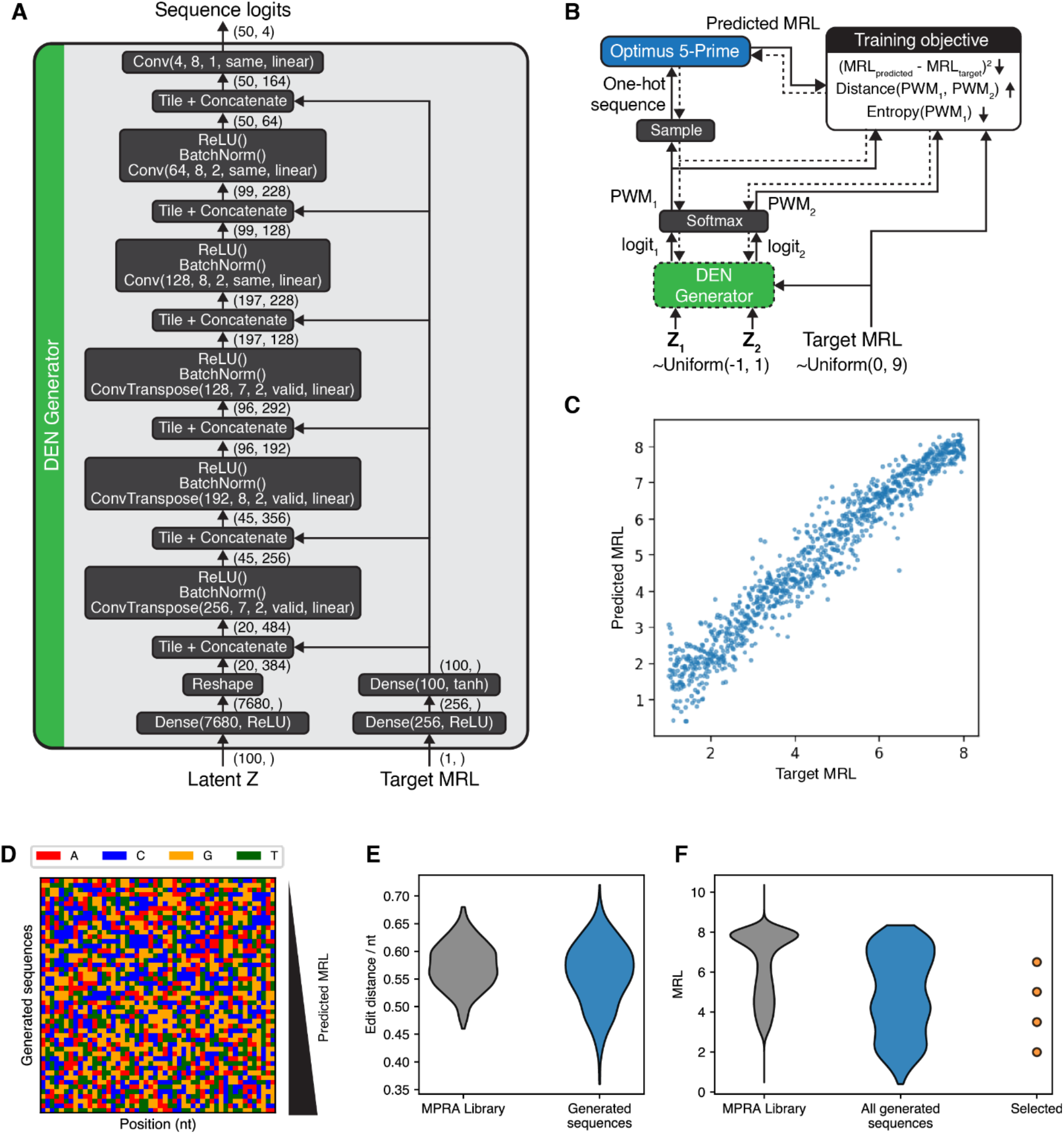
Design of 5’UTR sequences of varying MRLs using Optimus 5-Prime and an “inverse regression”-type Deep Exploration Network (DEN). **(A)** Architecture of the DEN generator network. Compared to **Supplementary Figure 7**, this network has an additional input representing the target MRL of the 5’UTR to be generated. Additionally, after being processed by two dense layers, this input is concatenated to the input of each convolutional layer. **(B)** DEN training schematic. Compared to **Supplementary Figure 7**, we additionally sample target MRL values from a uniformly random distribution during training, and set the fitness loss to the squared difference between the target and predicted MRL. Weights for the fitness, similarity, and entropy components of the loss function were 0.2, 5, and 1. Margin values for the similarity and entropy terms were 0.3 and 1.8. Training was performed for 100 epochs. **(C)** After DEN training, 1,024 5’UTR sequences covering a range of target MRLs were generated and compared to the MRL predicted by Optimus 5-Prime. **(D)** Colormap representation of 50 5’UTR sequences randomly chosen from the 1,024 generated by the trained DEN, sorted by predicted MRL. Each row represents a separate sequence, and color indicates nucleotide identity. **(E)** Distribution of edit distances per nucleotide, for 500 random pairs chosen from the MPRA library (left) or the 1,024 sequences generated by the DEN. **(F)** Distribution of MRLs measured from the MPRA library (left), or predicted by Optimus 5-Prime on all 1,024 DEN-generated sequences (middle) and the four selected sequences from this set (right).

**Supplementary Figure 9.**
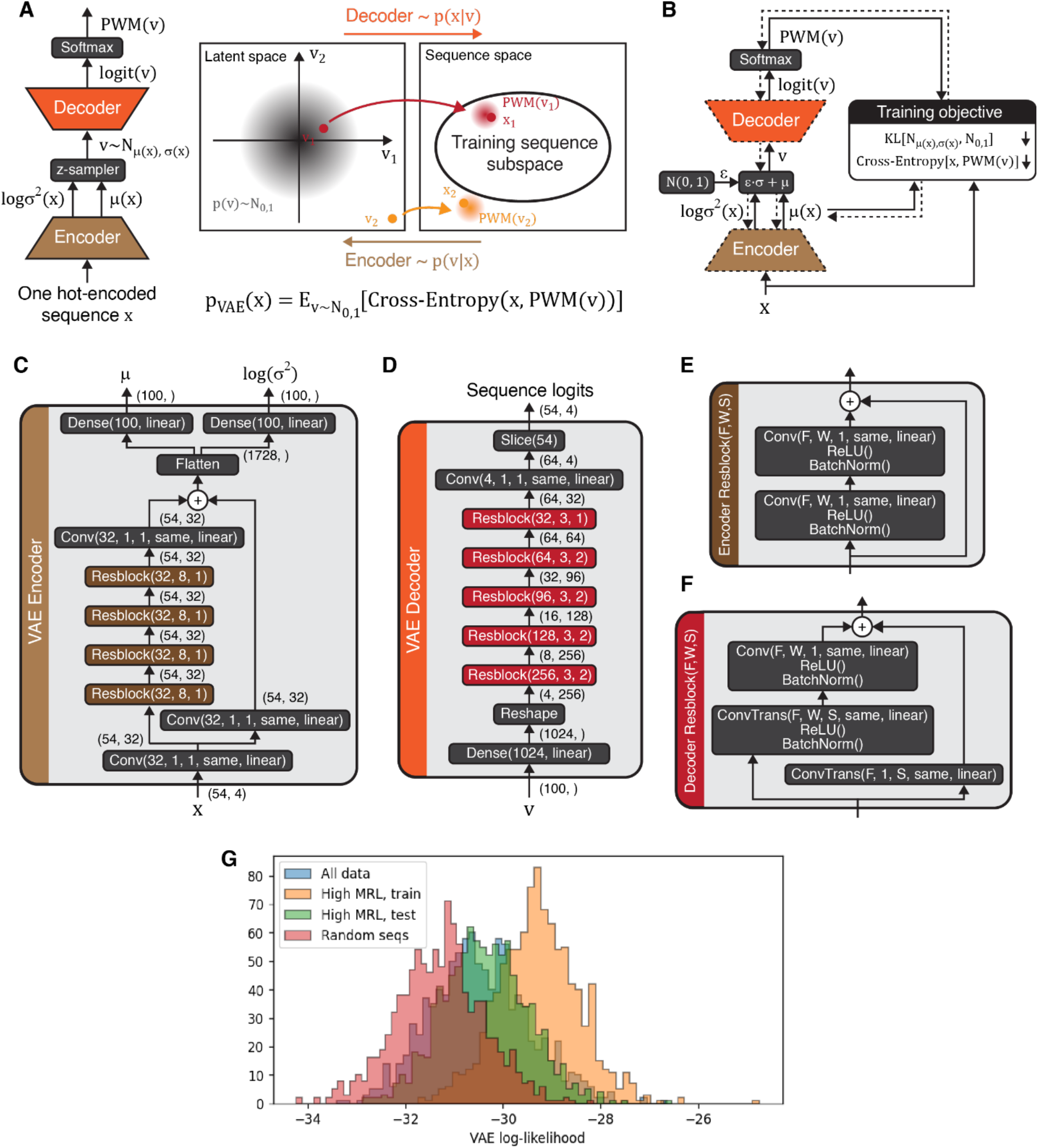
Variational Autoencoder (VAE) to estimate the likelihood of 5’UTR sequences in the fixed-end 50nt MPRA. **(A)** Schematic of a VAE structure and function. In the VAE framework, a sequence *x* originates from sampling a latent vector *v* from a continuous prior distribution *p*(*v*)∼*N*(0, 1), followed by sampling *x* from the likelihood *p*(*x*|*v*). Sequences in the dataset used for VAE training correspond to latent vectors close to 0 and therefore more likely under the prior. Two neural networks allow (probabilistic) conversion between the sequence and the latent space. On one hand, an encoder accepts a one hot-encoded *x* and returns the mean μ and log variance log(σ^2^) of a normal distribution corresponding to *p*(*v*|*x*). Conversely, a decoder converts a latent vector into a sequence logit, which can be converted into a PWM encoding *p*(*x*|*v*). Conceptually, the marginal probability of a sequence *p*(*x*) ≅ *p_VAE_*(*x*) is the expected cross-entropy (“distance”) between the *x* and the output *PWM*(*v*), when *v* is sampled from *p*(*v*) = *N*(0,1). In practice, it is more efficient to sample *v* from *N*(μ(*x*), σ^2^(*x*)) and use a correction factor to account for the different distribution (importance sampling). For implementation details, see^55^. **(B)** During VAE training, encoder and decoder weights are updated via gradient descent to minimize the KL-divergence between *N*(0,1) and *N*(μ(*x*), σ^2^(*x*)), as well as the cross-entropy between *x* and *PWM*(*v*), for all *x* in the training set. For a more comprehensive description of VAE training, see ^55^. **(C-F)** Architecture of the Encoder network **(C)**, decoder network **(D)**, and the residual blocks used in the encoder **(E)** and decoder **(F)**. Convolutional and transpose convolutional layers are represented as *Conv(F,W,S,P,A)* and *ConvTranspose(F,W,S,P,A)*, where *F* and *W* are the number and size of convolutional filters, *S* is the stride, *P* is the padding, and *A* is the activation function. For the 50nt VAE, a 54nt-long sequence is used where the last 4 bases are masked out with zeros. **(G)** Likelihood of different sequence sets under a VAE trained on high MRL 5’UTR sequences. Histograms were generated from 1,000 sequences randomly selected from the full MPRA library, the VAE training and testing sets, or randomly generated *in silico*.

**Supplementary Figure 10.**
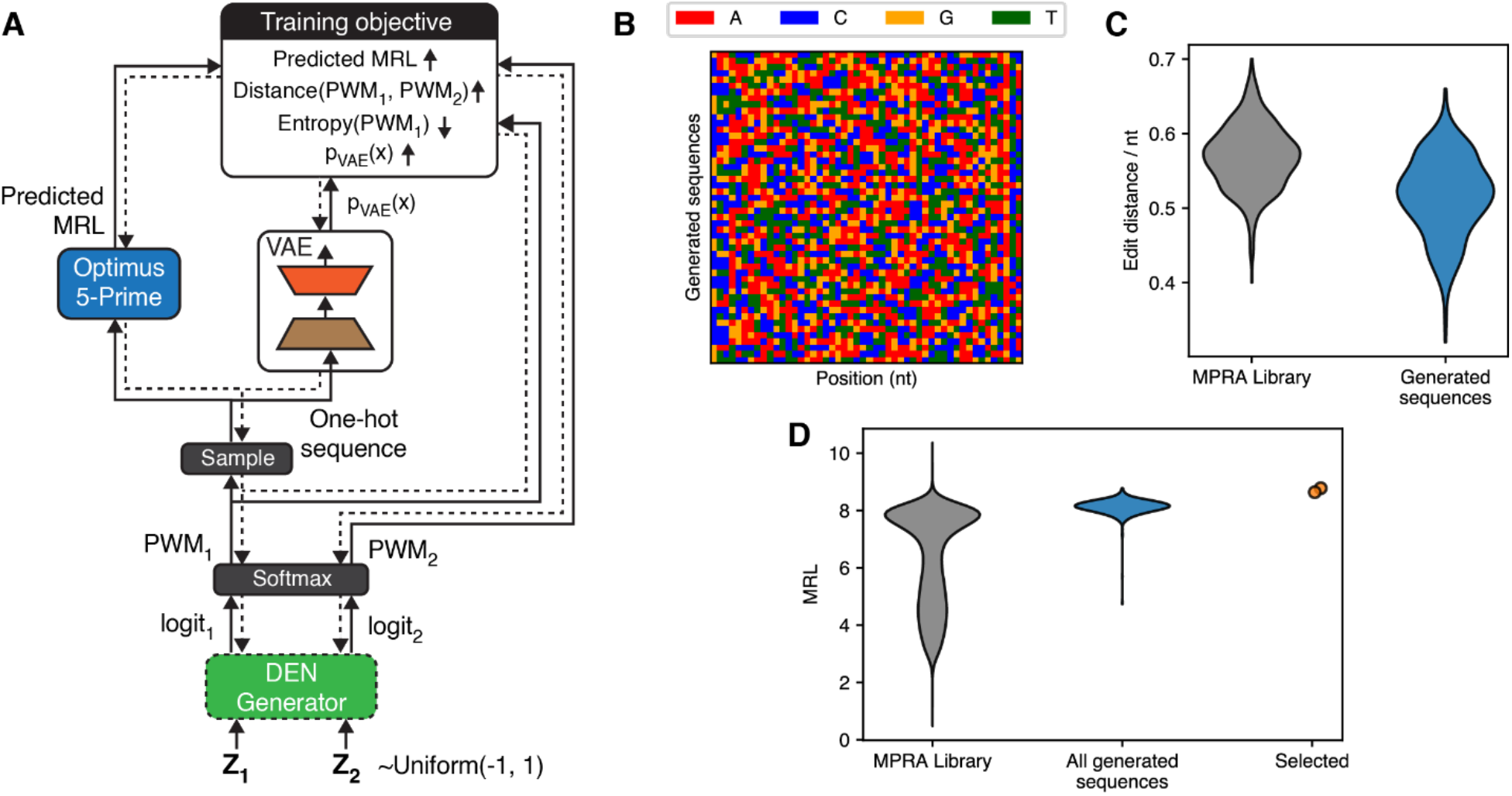
50nt 5’UTR design using Optimus 5-Prime, a Deep Exploration Network (DEN), and VAE regularization. **(A)** DEN training schematic. Compared to **Supplementary Figure 7**, we here used a VAE pretrained as shown in **Supplementary Figure 9** to estimate the marginal *p_VAE_*(*x*) of a generated sequence *x*, and we add a VAE component to the loss function set to max(0, *margin_VAE_* – *log*(*p_VAE_*(*x*))). Weights for the fitness, similarity, entropy, and VAE components of the loss function were 0.1, 5, 1, and 0.5. Margin values for the similarity, entropy, and VAE terms were 0.3, 1.8, and -30. Training was performed for 100 epochs. **(B)** Colormap representation of 50 5’UTR sequences randomly chosen from the 1,024 generated by the trained DEN. Each row represents a separate sequence, and color indicates nucleotide identity. **(C)** Distribution of edit distances per nucleotide, for 500 random pairs chosen from the MPRA library (left) or the 1,024 sequences generated by the DEN. **(D)** Distribution of MRLs measured from the MPRA library (left), or predicted by Optimus 5-Prime on all 1,024 DEN-generated sequences (middle) and the two selected sequences from this set (right).

**Supplementary Figure 11.**
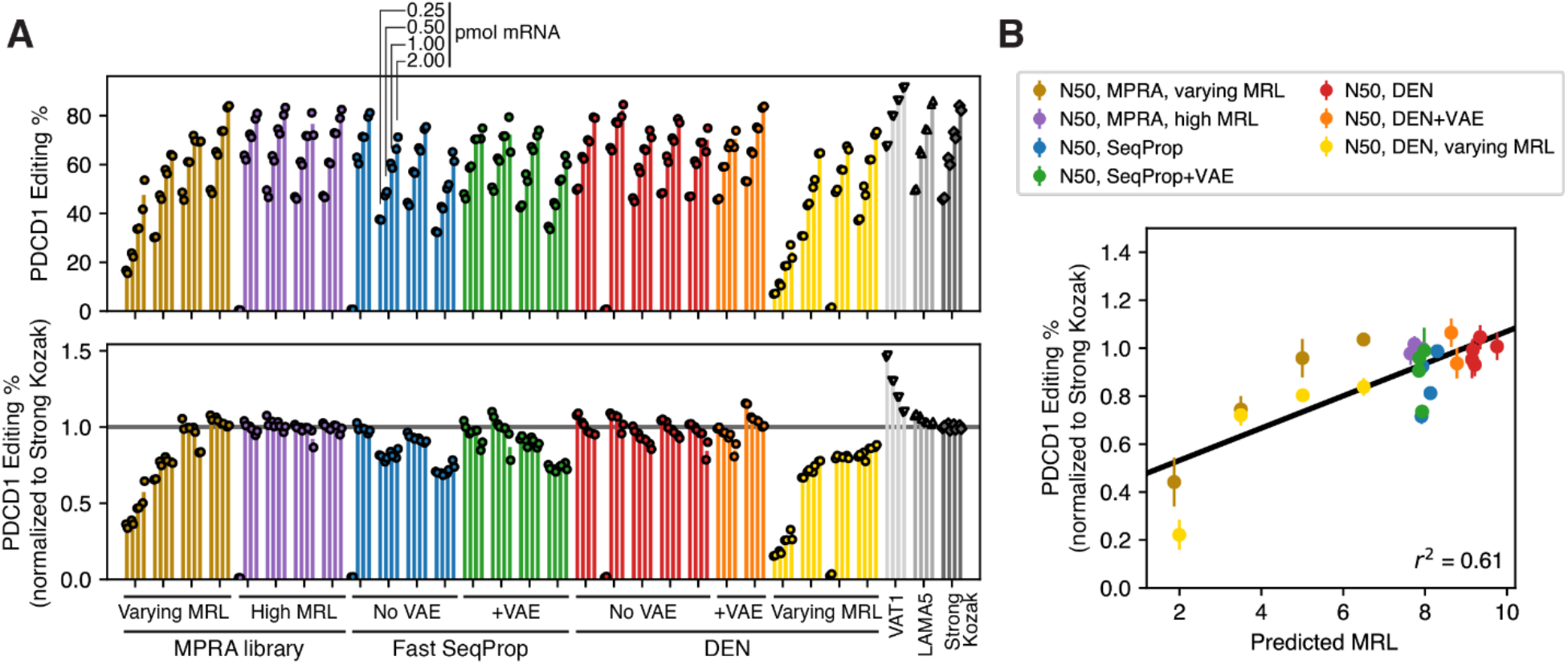
Performance of 50nt 5’UTR designs on the PDCD1 megaTAL. **(A)** Editing efficiencies for mRNAs with a megaTAL targeting the PDCD1 gene, for 30 different 5’UTR including designs and controls. Top: absolute editing efficiencies. Bottom: editing efficiencies normalized to the Strong Kozak control. Analogous to Figure 2C but with the PDCD1 megaTAL instead of TGFBR2. Editing efficiencies for the first High MRL MPRA control, the first No VAE Fast SeqProp design, the second No VAE DEN design, and the third Varying MRL DEN design were close to zero only at a dosage of 0.25 pmol mRNA, and were deemed to be the result of experimental error and excluded from subsequent analysis. **(B)** Kozak-normalized editing efficiency of the PDCD1 megaTAL vs. Optimus 5-Prime predicted MRL for all designed and MPRA control 5’UTRs. Analogous to Figure 2E but with the PDCD1 megaTAL instead of TGFBR2.

**Supplementary Figure 12.**
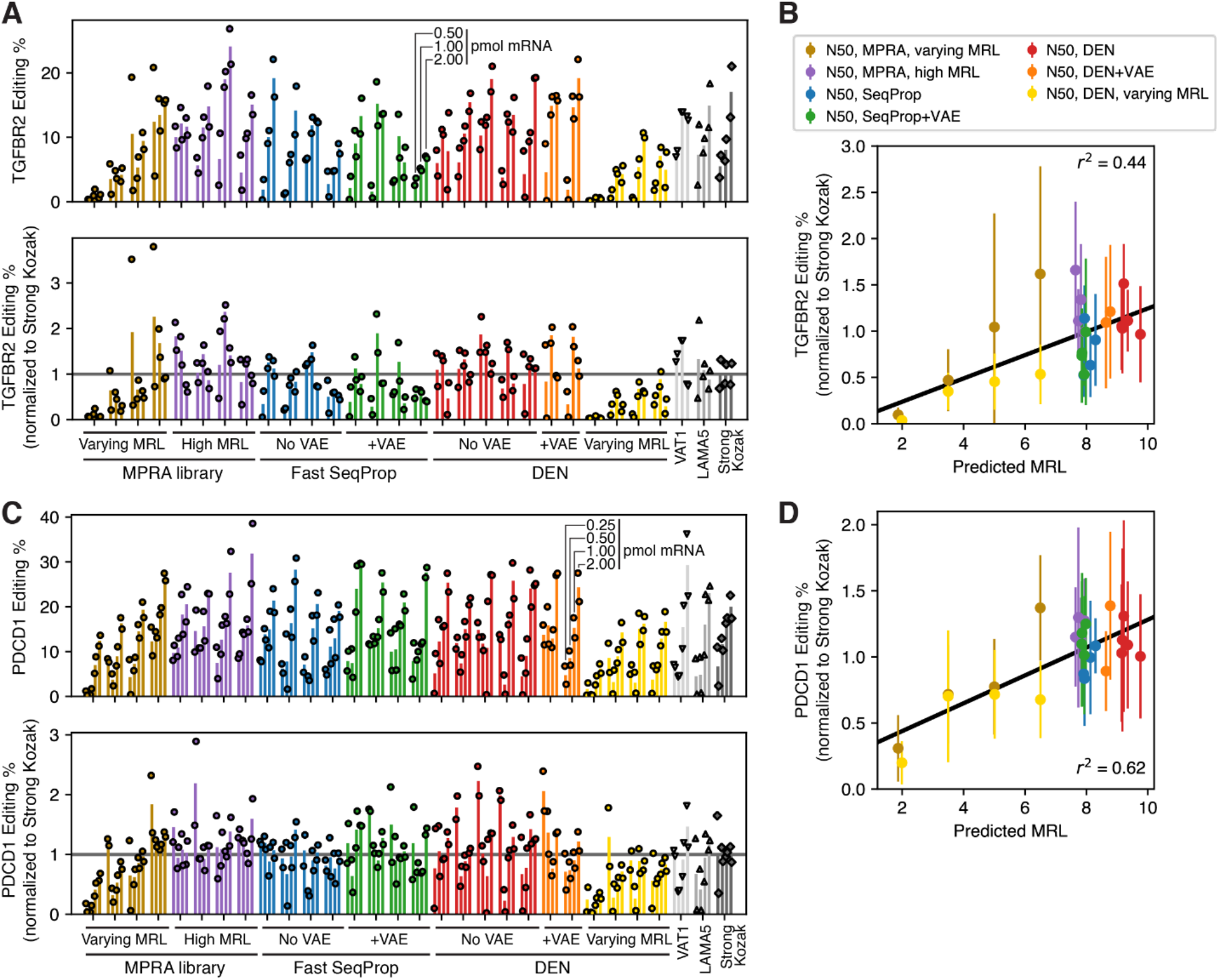
Performance of 50nt 5’UTR designs on the TGFBR2 and PDCD1 megaTALs in HepG2. **(A and C)** Editing efficiencies for mRNAs with a megaTAL targeting the TGFBR2 **(A)** or PDCD1 **(C)** genes in HepG2 cells, for 30 different 5’UTR including designs and controls. Top: absolute editing efficiencies. Bottom: editing efficiencies normalized to the Strong Kozak control. Analogous to Figure 2C and **Supplementary Figure 11A** but using HepG2 cells instead of K562. Only three mRNA dosage levels were evaluated for TGFBR2. **(B and D)** Kozak-normalized editing efficiencies of the TGFBR2 **(B)** and PDCD1 **(D)** megaTALs vs. Optimus 5-Prime predicted MRL for all designed and MPRA control 5’UTRs. Analogous to Figure 2E and **Supplementary Figure 11B** but using HepG2 cells instead of K562.

**Supplementary Figure 13.**
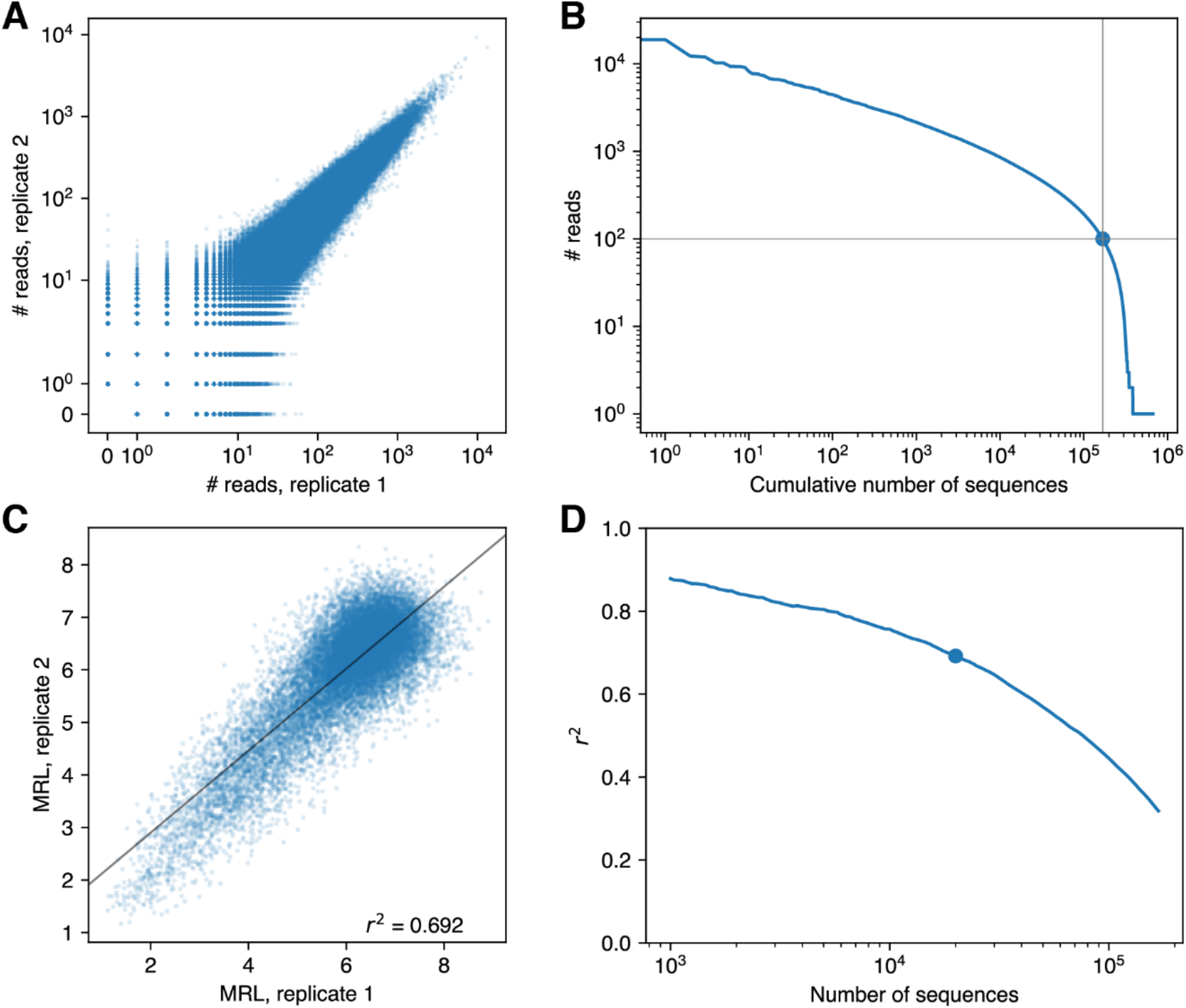
Basic analysis of random-end N25 MPRA library. **(A)** Sequencing read coverage for all sequences across two biological replicates. **(B)** Number of sequences resulting from a given cutoff on the total number of reads per sequence across both replicates. Marker indicates 168,297 sequences with at least 100 reads. **(C)** MRL correlation across replicates, for the top 20,000 sequences by read coverage. Black line represents a regression line. **(D)** r^2^ as a function of the number of sequences used. Sequences were sorted by the total number of reads across both replicates in descending order. Then, the top x sequences (x axis) were used to calculate a corresponding r^2^ value (y axis). The large marker indicates the number of sequences and r^2^ in **(C)**.

**Supplementary Figure 14.**
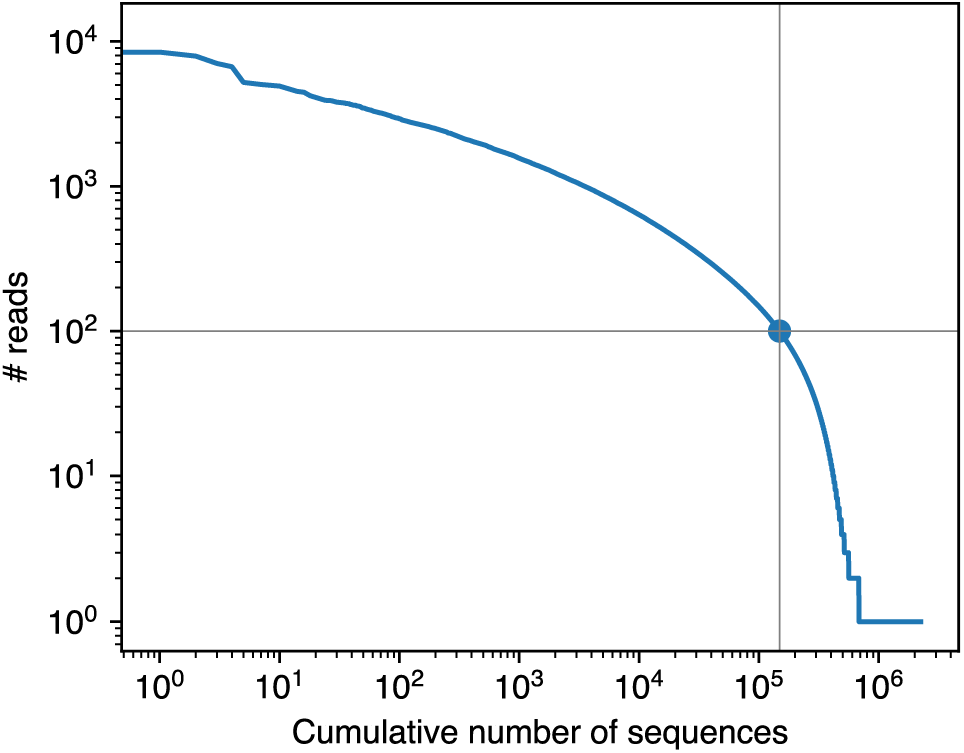
Read coverage as a function of the number of sequences retained in the random-end N50 MPRA. Marker indicates 57,165 sequences with at least 100 reads.

**Supplementary Figure 15.**
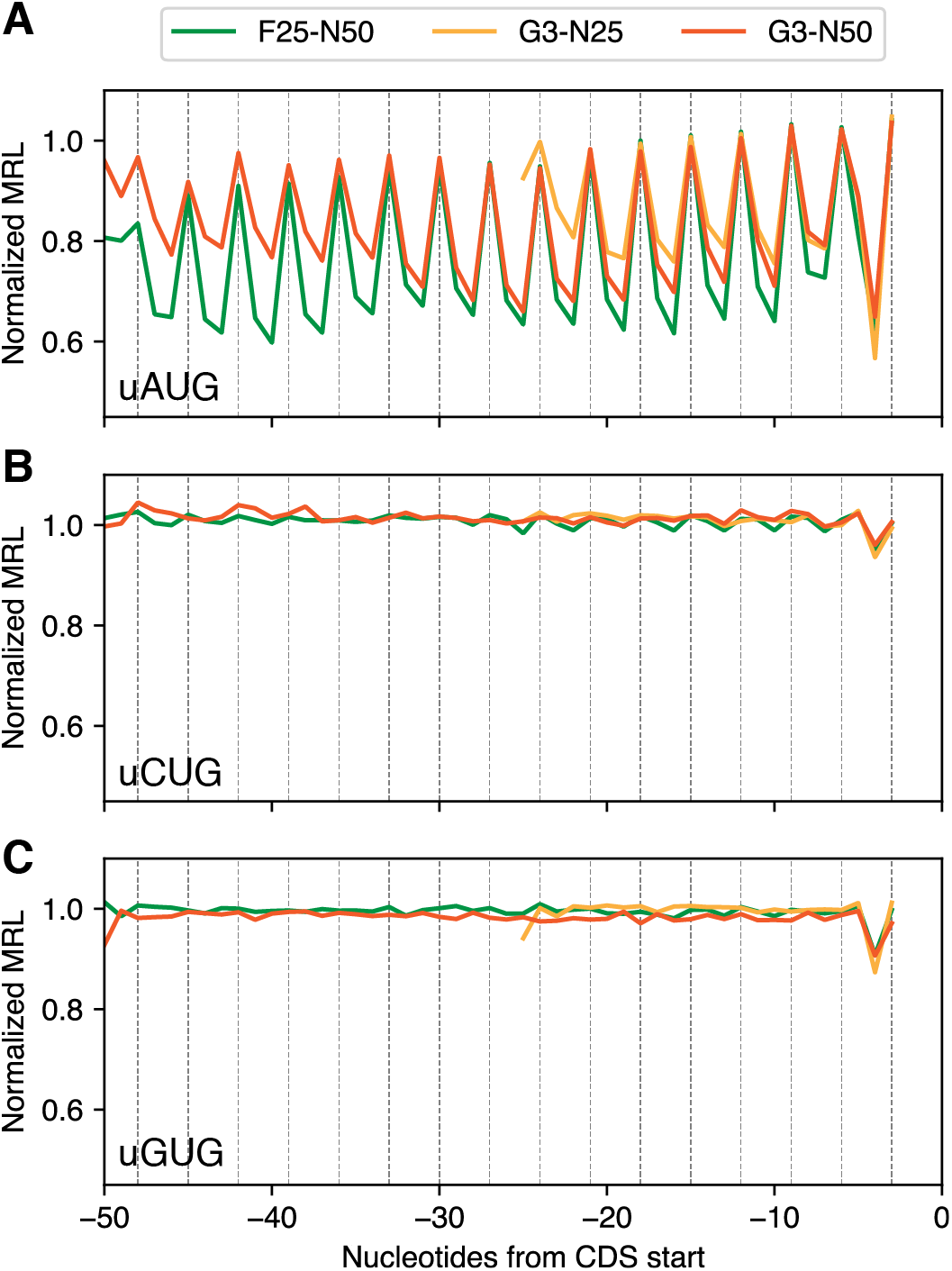
Effects of upstream start codons on MRL in three MPRA libraries. Median MRL of all sequences containing a uAUG **(A)**, uCUG **(B)**, and uGUG **(C)** at the indicated position from the start of the EGFP ORF, for the 25nt-(yellow) or 50nt-long (orange) randomized 5’UTR libraries, as well as our previous “fixed-end” 50nt library (green). MRL was normalized to the median of each library. **(A)** is identical to Figure 3B but aligned to the EGFP start codon instead of the start of the transcript.

**Supplementary Figure 16.**
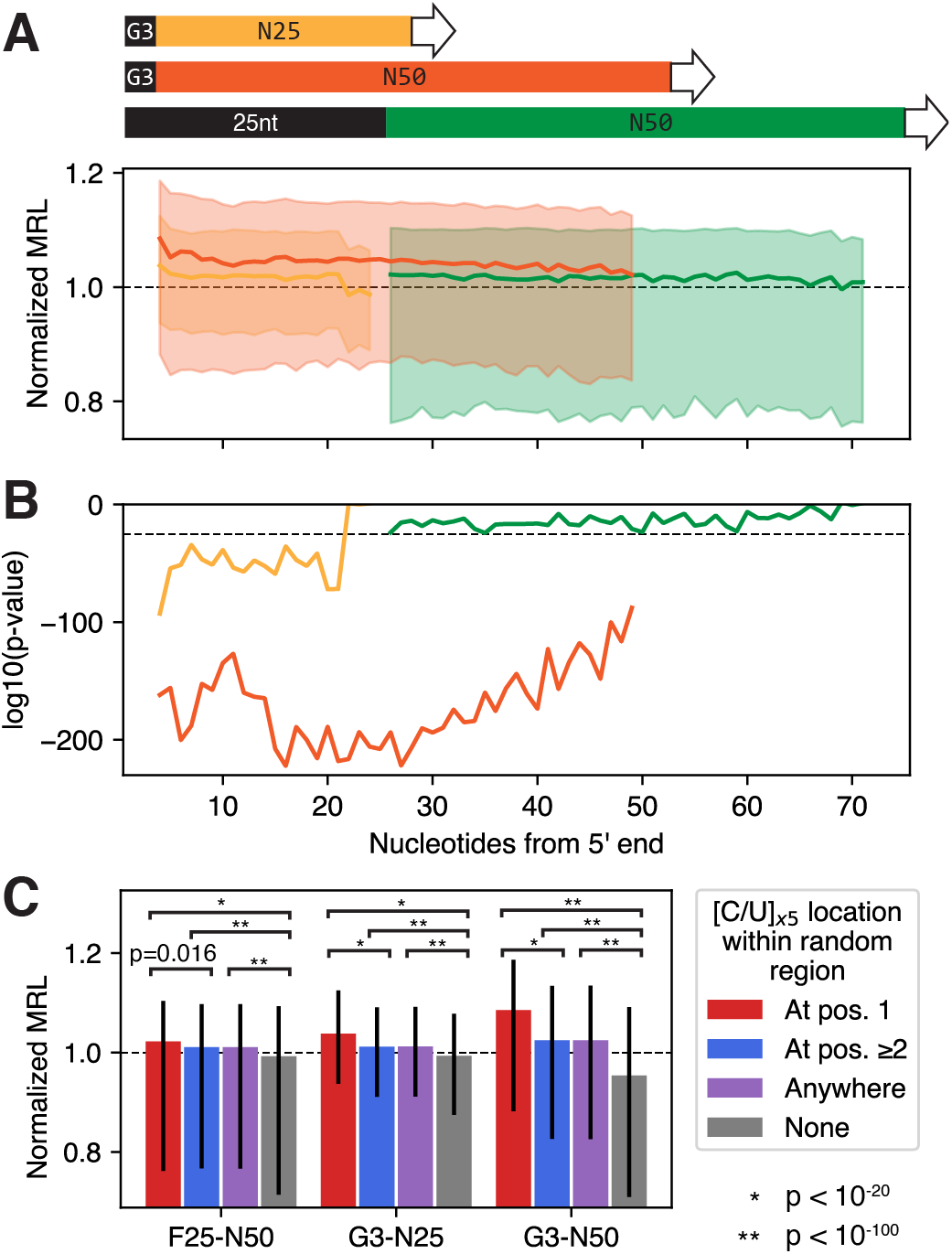
Detailed analysis of the effects of 5’UTR polypyrimidine tracts on MRL in three MPRA libraries. **(A)** Median MRL (solid lines) and interquartile range (shaded regions) of all sequences containing 5nt-long polypyrimidine (C or U) tract at the indicated position from the start of the transcript. Identical to Figure 3C but with the interquartile range indicated. **(B)** Bonferroni-corrected p-values from a Mann Whitney U test of medians between the MRL of sequences that contain a 5nt-long polypyrimidine tract at the indicated position versus sequences that do not contain 5nt-long polypyrimidine tracts at all. Horizontal bar indicates p = 10^-25^. **(C)** MRL of library sequences containing a 5nt-long polypyrimidine tract (C or U) within the random region, starting at position 1 (red bars), 2 or after (blue bars), anywhere (purple bars), or none at all (gray bars). MRLs were normalized to the median of each library as in **(A)**. Bars indicate the normalized median MRL within each group. Error bars indicate the interquartile range. Horizontal brackets indicate Bonferroni-corrected p-values from a Mann Whitney U test of medians between the MRLs in each group. *: p < 10^-20^, **: p < 10^-100^.

**Supplementary Figure 17.**
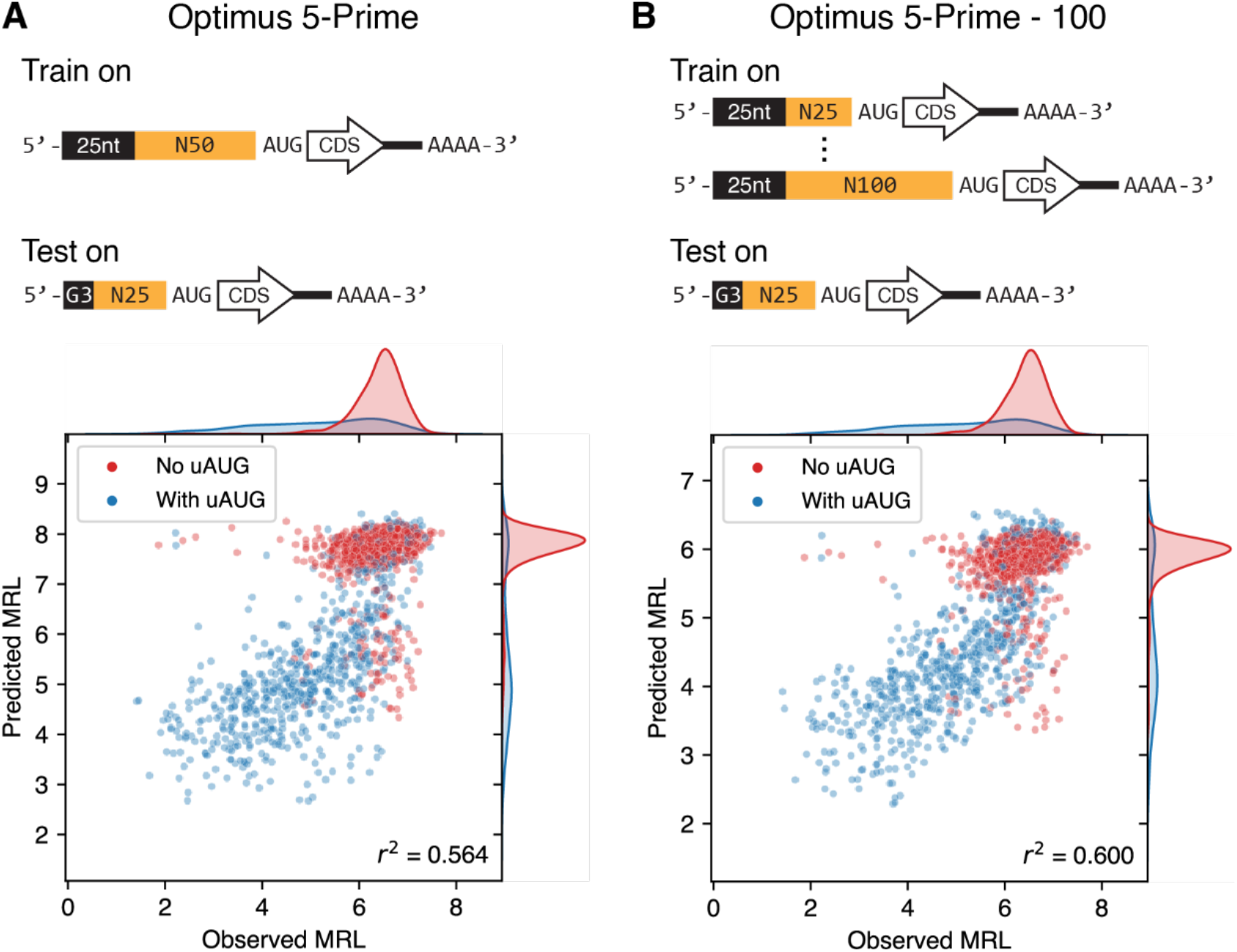
Performance of previously developed Optimus 5-Prime models on the random-end 25nt library. **(A)** Performance of the original Optimus 5-Prime trained on the fixed-end 50nt 5’UTR library. **(B)** Performance of Optimus 5-Prime - 100^33^, a model with the same architecture as the original Optimus 5-Prime but trained on a 5’UTR library with a fixed 25nt segment followed by a variable region between 25 and 100nt long. As in Figure 3E, these models were evaluated against a test dataset comprised of the 2,000 sequences with the highest read coverage in the random-end 25nt MPRA library.

**Supplementary Figure 18.**
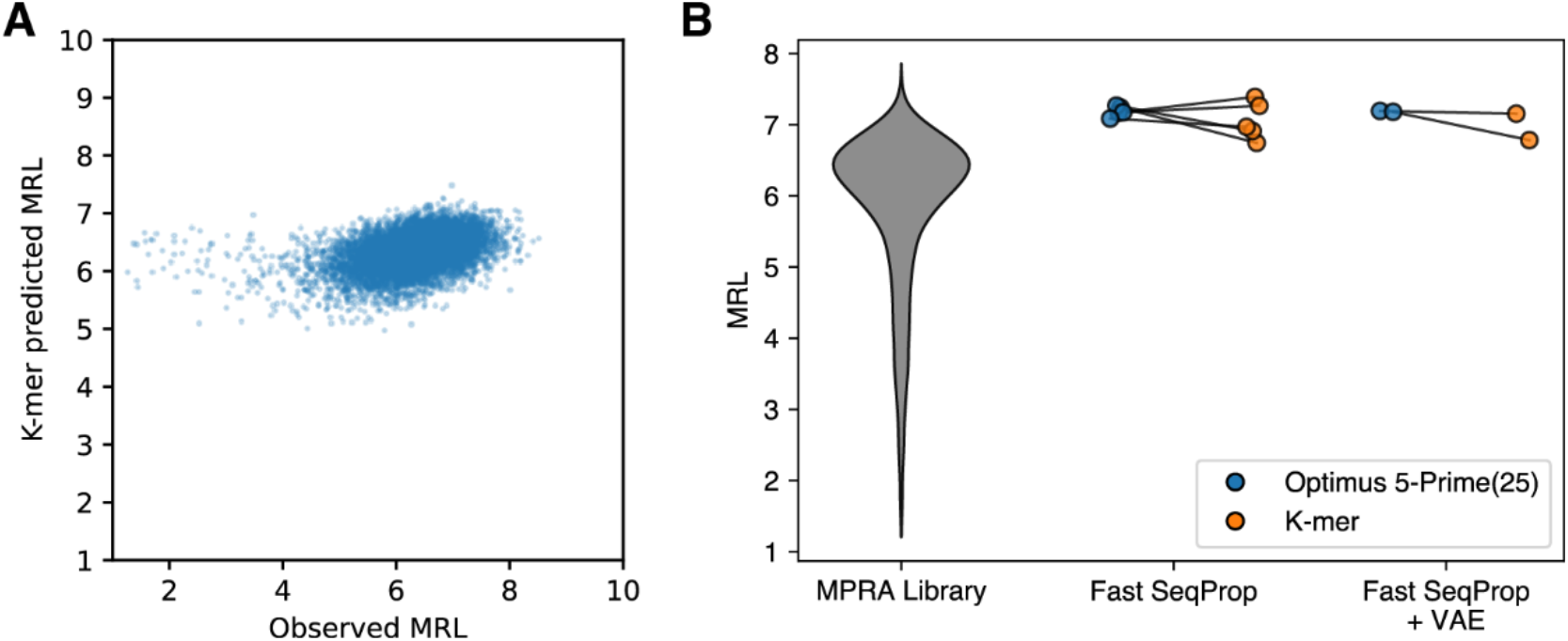
Using a linear k-mer model to validate 25nt-long Fast SeqProp designs. See **Methods** for training and model details. **(A)** Observed vs. predicted MRL on a held-out set of 11,349 sequences with no uAUG and greater than 200 reads. Pearson r = 0.4094. **(B)** Comparison of predicted MRL for the five sequences designed via Fast SeqProp and the two sequences designed via Fast SeqProp with VAE regularization, when using Optimus 5-Prime(25) or the k-mer predictors. A violin of the entire MPRA library is shown on the left for comparison. Compared to k-mer model predictions on the entire test set shown in **(A)**, most predictions on the designed sequences are within the top 1%. The exceptions are two sequences designed without VAE regularization, which are within the top 1.2% and 6.3%, and one VAE-regularized design, which is within the top 4.5%.

**Supplementary Figure 19.**
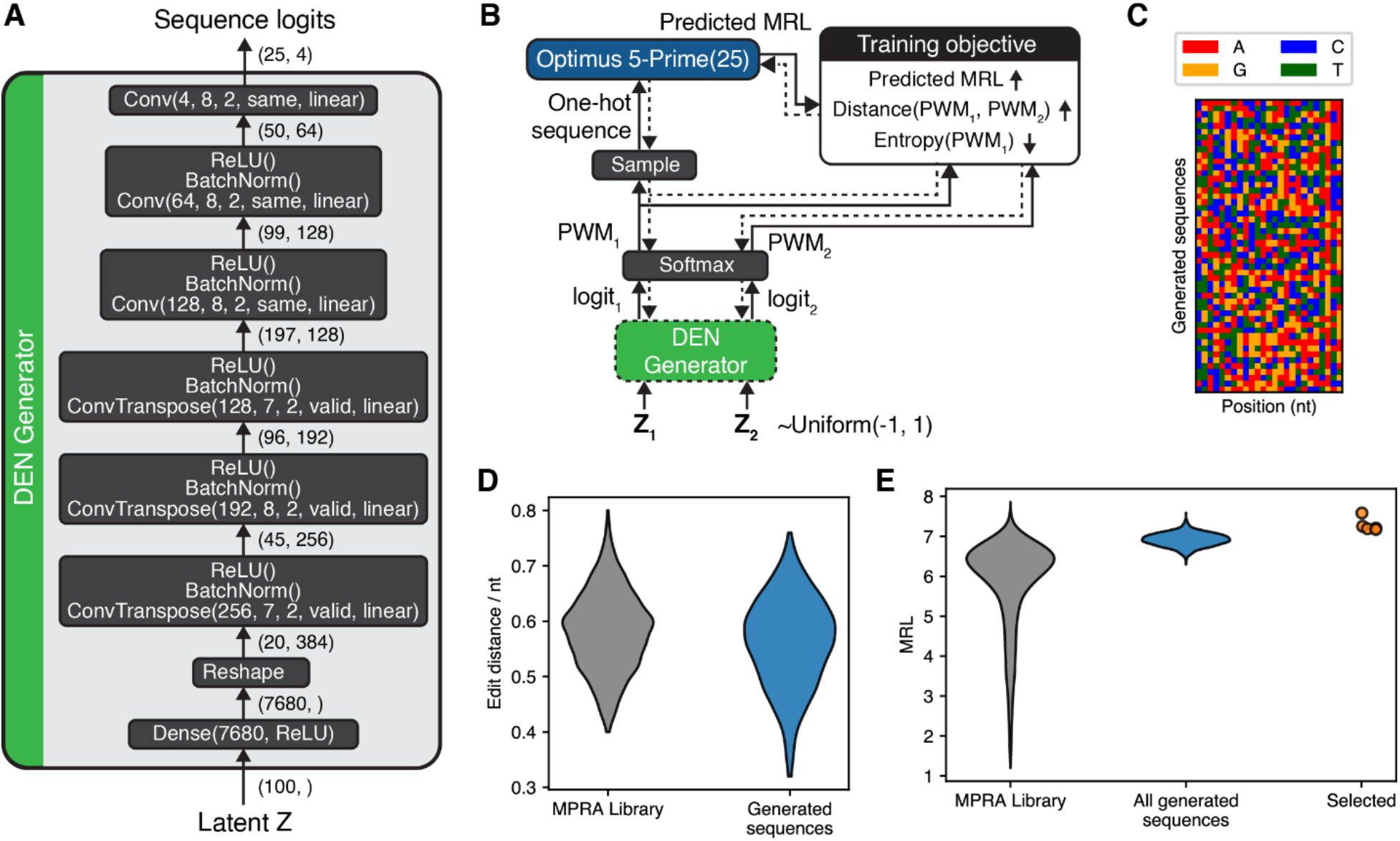
25nt 5’UTR design using Optimus 5-Prime(25) and Deep Exploration Networks. **(A)** Architecture of the DEN generator network, which takes a continuous-valued 100-dimensional latent vector and returns a 25×4-dimesional continuous-valued logit representing a sequence. Convolutional and Transpose convolutional layers are represented as *Conv(F,W,S,P,A)* and *ConvTranspose(F,W,S,P,A)*, where *F* and *W* are the number and size of convolutional filters, *S* is the stride, *P* is the padding, and *A* is the activation function. **(B)** DEN training schematic. Training was performed as described in **Supplementary Figure 7** but with Optimus 5-Prime(25) (Figure 3) as the predictor. Weights for the fitness, similarity, and entropy components of the loss function were 0.35, 5, and 1. Margin values for the similarity and entropy terms were 0.3 and 1.8. Training was performed for 100 epochs. **(C)** Colormap representation of 50 5’UTR sequences randomly chosen from the 1,024 generated by the trained DEN. Each row represents a separate sequence, and color indicates nucleotide identity. **(D)** Distribution of edit distances per nucleotide, for 500 random pairs chosen from the 25nt random-end MPRA library (left) or the 1,024 sequences generated by the DEN. **(E)** Distribution of MRLs measured from the MPRA library (left), or predicted by Optimus 5-Prime(25) on all 1,024 DEN-generated sequences (middle) and the four selected sequences from this set (right).

**Supplementary Figure 20.**
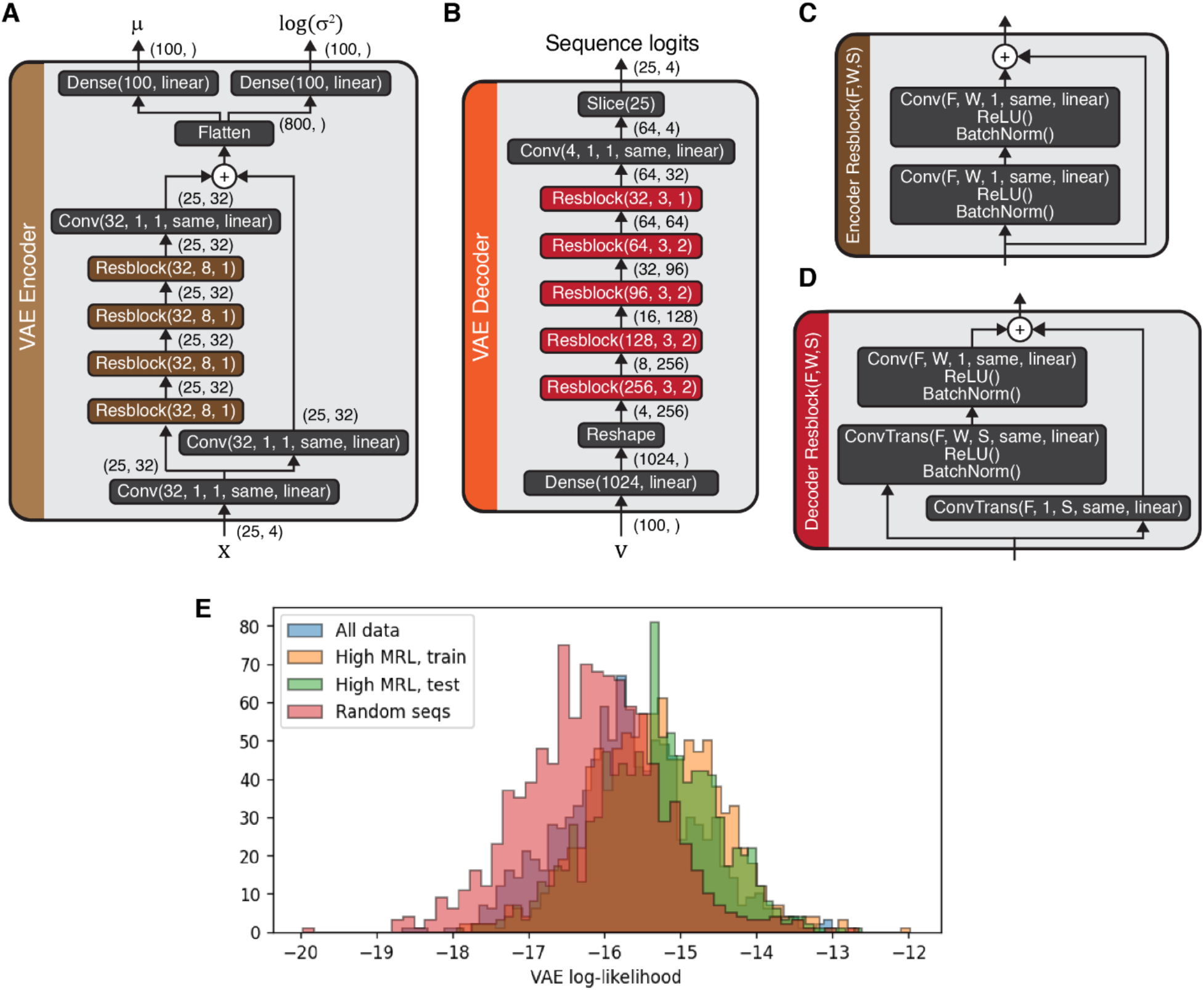
Variational Autoencoder (VAE) to estimate the likelihood of 5’UTR sequences in the 25nt random-end MPRA. The general VAE architecture and training scheme is identical to those in **Supplementary Figure 9A** and B. **(A-D)** Architecture of the Encoder network **(A)**, decoder network **(B)**, and the residual blocks used in the encoder **(C)** and decoder **(D)**. Convolutional and transpose convolutional layers are represented as *Conv(F,W,S,P,A)* and *ConvTranspose(F,W,S,P,A)*, where *F* and *W* are the number and size of convolutional filters, *S* is the stride, *P* is the padding, and *A* is the activation function. **(E)** Likelihood of different sequence sets under a VAE trained on high MRL 5’UTR sequences. Histograms were generated from 1,000 sequences randomly selected from the full MPRA library, the VAE training and testing sets, or randomly generated in silico.

**Supplementary Figure 21.**
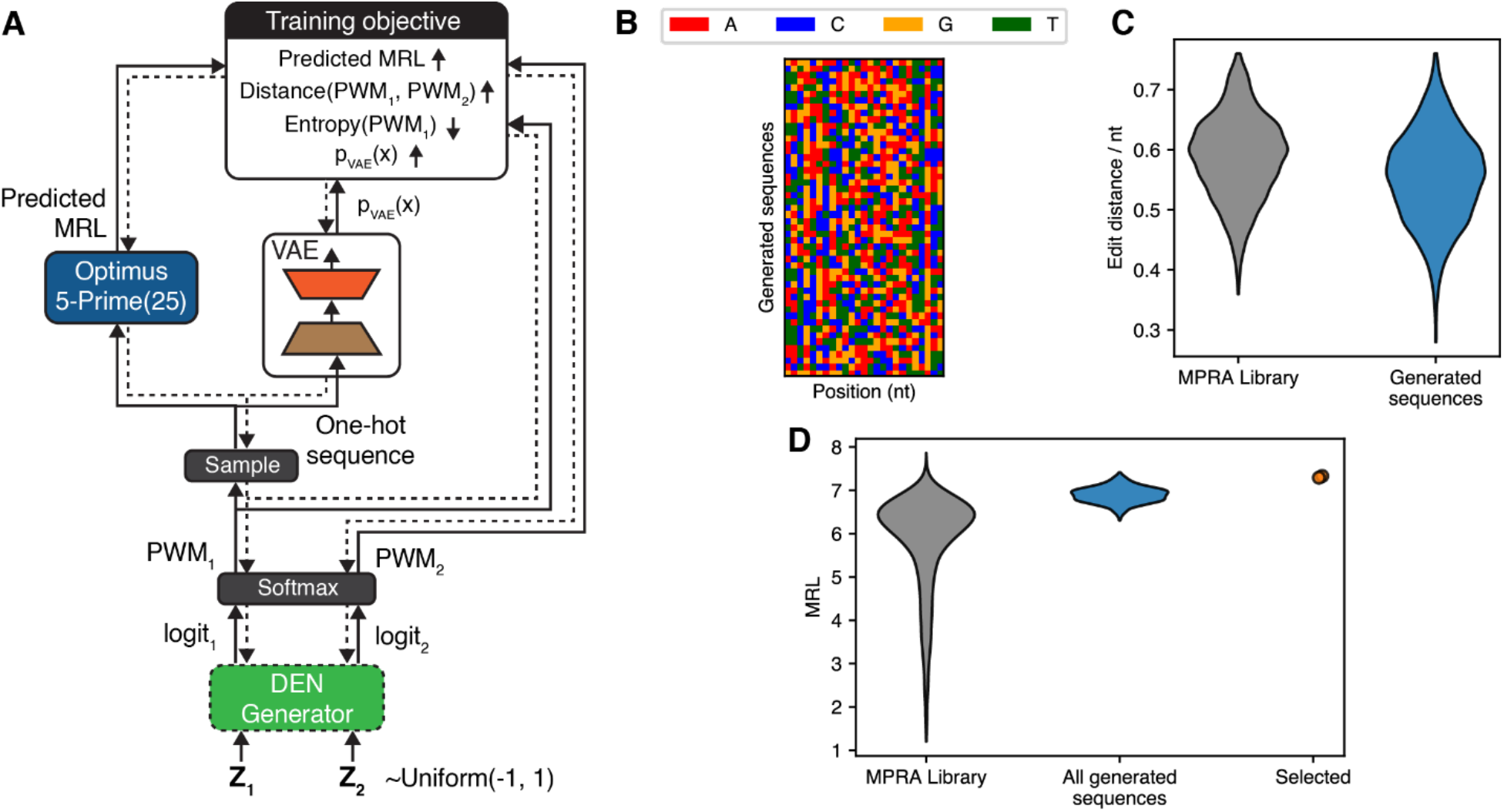
25nt-long 5’UTR design using Optimus 5-Prime(25), a Deep Exploration Network (DEN), and VAE regularization. **(A)** DEN training schematic. Compared to **Supplementary Figure 19**, we use a VAE, pretrained as shown in **Supplementary Figure 20**, to estimate the marginal *p_VAE_*(*x*) of a generated sequence *x*, and we add a VAE component to the loss function set to max(0, *margin_VAE_* – *log*(*p_VAE_*(*x*))). Weights for the fitness, similarity, entropy, and VAE components of the loss function were 0.3, 5, 1, and 0.5. Margin values for the similarity, entropy, and VAE terms were 0.3, 1.8, and -30. Training was performed for 100 epochs. **(B)** Colormap representation of 50 5’UTR sequences randomly chosen from the 1,024 generated by the trained DEN. Each row represents a separate sequence, and color indicates nucleotide identity. **(C)** Distribution of edit distances per nucleotide, for 500 random pairs chosen from the 25nt random-end MPRA library (left) or the 1,024 sequences generated by the DEN. **(D)** Distribution of MRLs measured from the MPRA library (left), or predicted by Optimus 5-Prime(25) on all 1,024 DEN-generated sequences (middle) and the two selected sequences from this set (right).

**Supplementary Figure 22.**
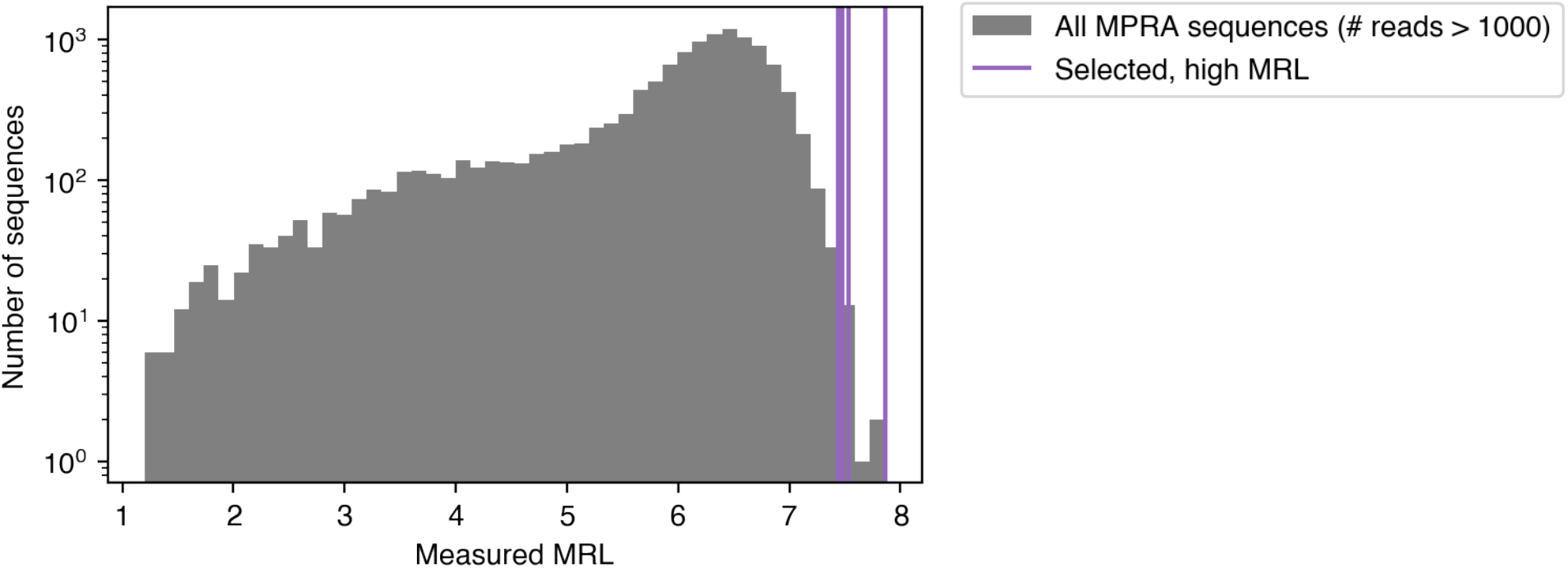
Random-end 25nt MPRA “high MRL” controls for the megaTAL gene editing assays. (A) Starting from the random-end 25nt MPRA library data in HEK293T, we excluded sequences if their read count was lower than 1,000 or if they contained uATGs. The remaining sequences were sorted by MRL, and four from the top twenty were selected. The histogram shows the measured MRLs of all four control sequences, compared with a high-coverage (# reads > 1,000) subset of the MPRA library.

**Supplementary Figure 23.**
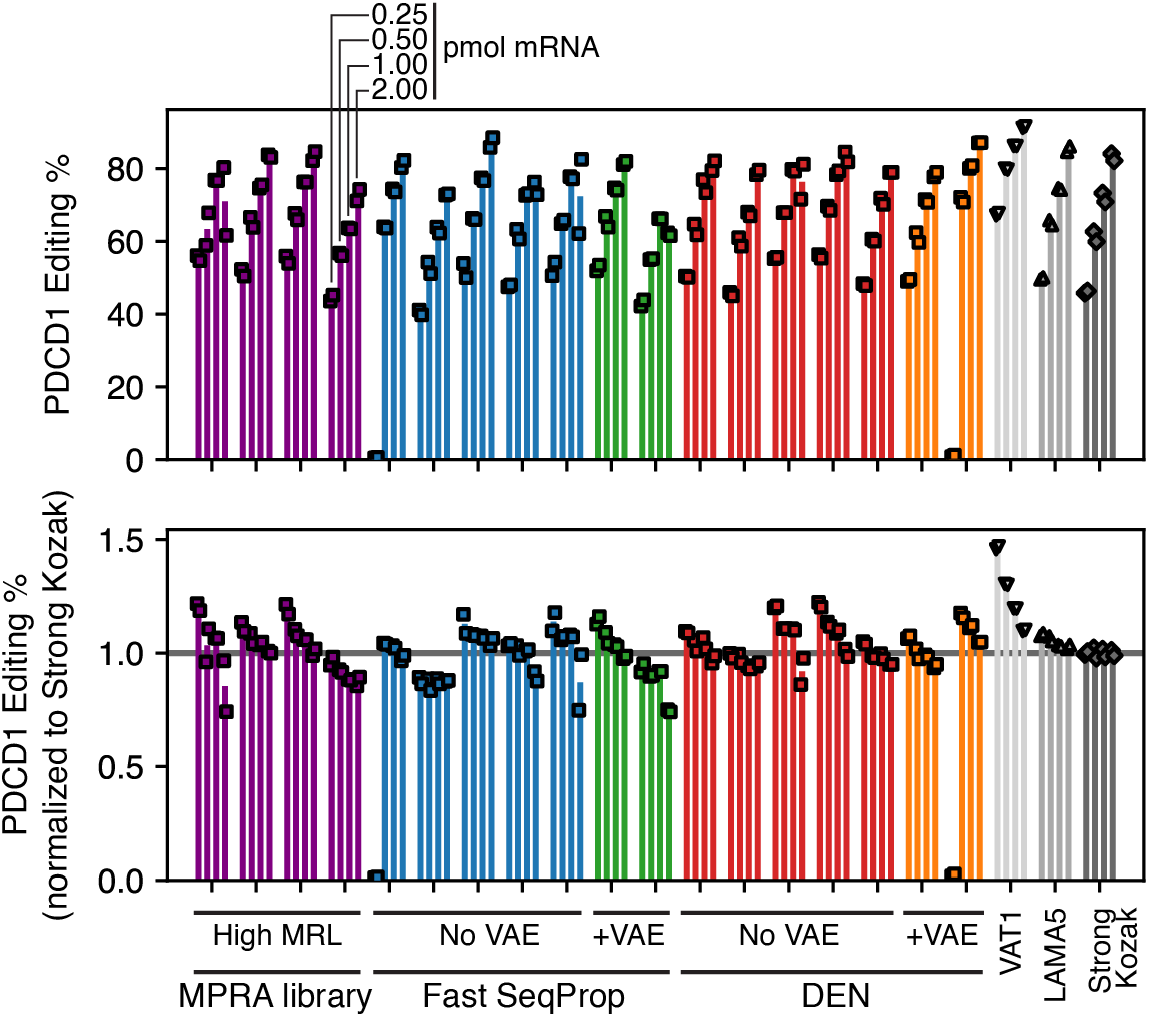
Performance of the 25nt 5’UTR designs on the PDCD1 megaTAL. Editing efficiencies for mRNAs with a megaTAL targeting the PDCD1 gene, for 21 different 5’UTRs including designs and controls. Top: absolute editing efficiencies. Bottom: editing efficiencies normalized to the Strong Kozak control. Analogous to Figure 4A and B but with the PDCD1 megaTAL instead of TGFBR2.

**Supplementary Figure 24.**
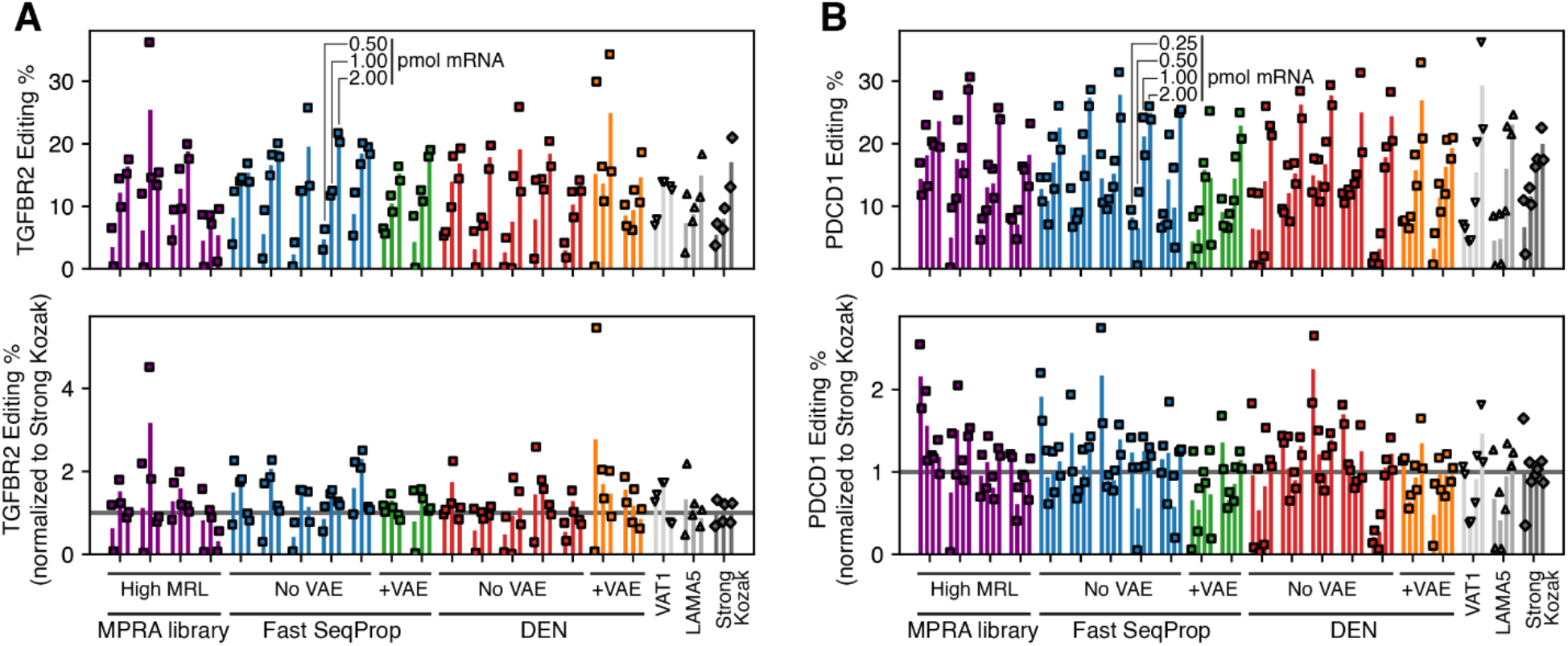
Performance of 25nt 5’UTR designs on TGFBR2 and PDCD1 megaTALs in HepG2. Editing efficiencies for mRNAs with a megaTAL targeting the TGFBR2 **(A)** or PDCD1 **(B)** genes in HepG2 cells, for 21 different 5’UTRs including designs and controls. Top: absolute editing efficiencies. Bottom: editing efficiencies normalized to the Strong Kozak control. Analogous to Figure 4A-B and **Supplementary Figure 23** but using HepG2 instead of K562.

**Supplementary Figure 25.**
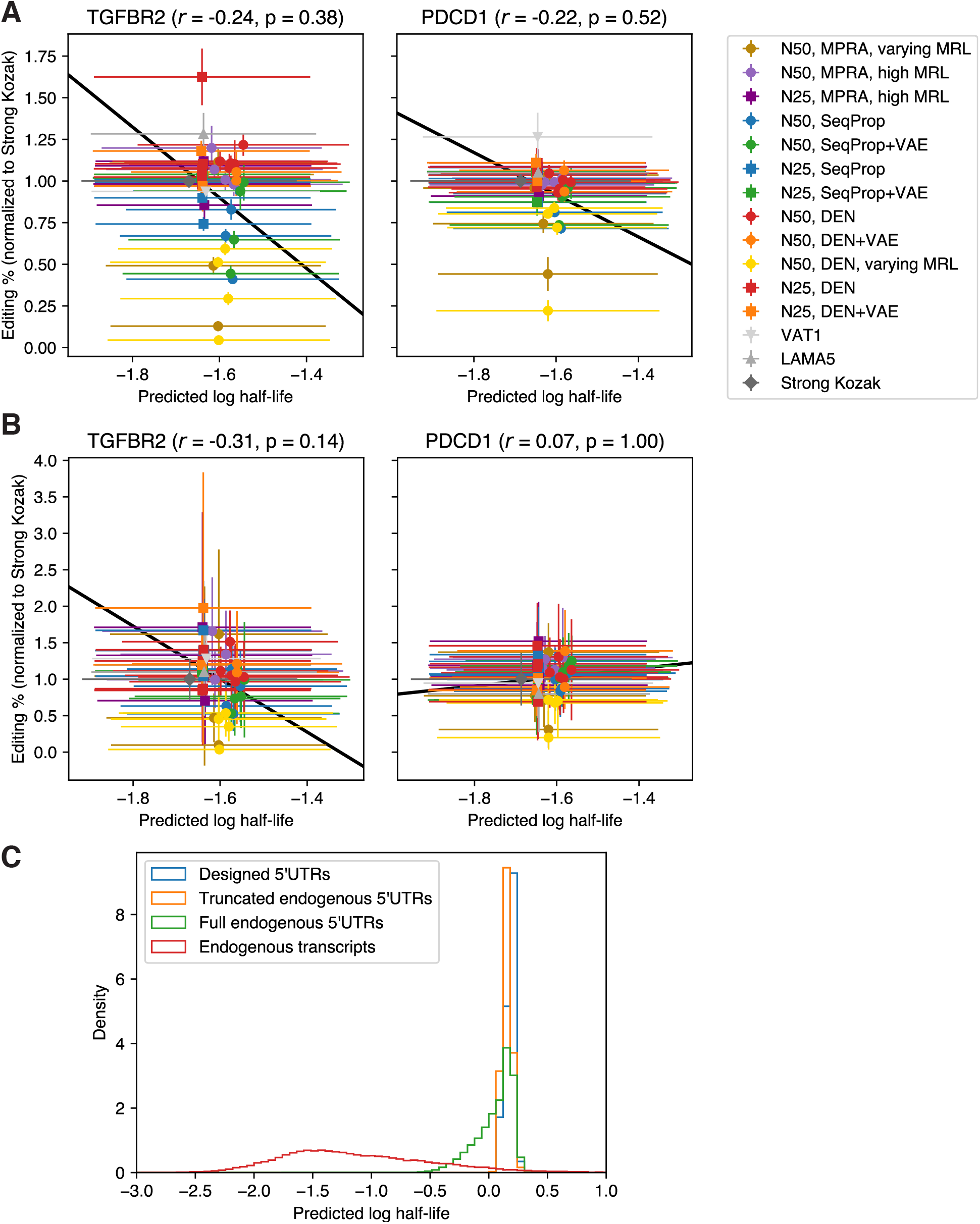
Half-life predictions for mRNAs containing all designed 5’UTRs versus editing efficiencies. Predictions were made using Saluki^35^, a model comprised of an ensemble of 50 hybrid convolutional/recurrent neural networks. Inputs to the predictor are a one-hot encoded mRNA sequence, a binary sequence indicating whether each base corresponds to the first base of a codon in the main ORF, and another binary sequence indicating splice sites. Outputs are log half-life predictions for each model in the ensemble. For this analysis, the splice site sequence was set to all zeros to reflect the absence of splicing in IVT mRNA. **(A and B)** Kozak-normalized editing efficiencies in K562 **(A)** and HepG2 **(B)** versus predicted log half-life. X coordinates of markers and horizontal error bars are the mean and standard deviations of Saluki ensemble model predictions. Titles indicate whether editing efficiencies correspond to TGFBR2 or PDCD1 megaTALs, and their respective Pearson correlation coefficients. **(C)** 5’UTRs have a limited ability to tune Saluki-predicted mRNA stability. Here, we used the EGFP ORF and the BGH-derived 3’UTR to mimic the conditions in our MPRA assay. “Designed 5’UTRs” correspond to those in **(A)** and **(B)**. “Endogenous transcripts” correspond to all protein-coding full transcripts from ensembl without splicing site annotations. “Full endogenous 5’UTRs” correspond to the same endogenous ensembl 5’UTRs along with the EGFP CDS and BGH 3’UTR. In “Truncated endogenous 5’UTRs”, the final 50 nt of every endogenous 5’UTR was placed downstream of the constant 25 nt 5’UTR sequence from our MPRA library, along with the EGFP CDS and BGH 3’UTR.

